# Sodium channel subpopulations with distinct biophysical properties and subcellular localization enhance cardiac conduction

**DOI:** 10.1101/2023.03.09.531903

**Authors:** Seth H. Weinberg

**Affiliations:** Department of Biomedical Engineering, The Ohio State University

## Abstract

Sodium (Na^+^) current is responsible for the rapid depolarization of cardiac myocytes that triggers the cardiac action potential upstroke. Recent studies have illustrated the presence of multiple “pools” of Na^+^ channels with distinct biophysical properties and subcellular localization, including clustering of channels at the intercalated disk and along the lateral membrane. Computational studies predict that Na^+^ channel clusters at the intercalated disk can regulate cardiac conduction via modulation of the narrow intercellular cleft between electrically coupled myocytes. However, these studies have primarily focused on the redistribution of Na^+^ channels between intercalated disk and lateral membranes and not considered the distinct biophysical properties of the Na^+^ channel subpopulations. In this study, we simulate computational models of single cardiac cells and one-dimensional cardiac tissues to predict the functional consequence of distinct Na^+^ channel subpopulations. Single cell simulations predict that a subpopulation of Na^+^ channels with shifted steady-state activation and inactivation voltage-dependency promote an earlier action potential upstroke. In cardiac tissues that account for distinct subcellular spatial localization, simulations predict that “shifted” Na^+^ channels contribute to faster and more robust conduction, with regards to sensitivity to tissue structural changes (i.e., cleft width), gap junctional coupling, and rapid pacing rates. Simulations predict that the intercalated disk-localized shifted Na^+^ channels provide a disproportionally larger contribution to total Na^+^ charge, relative to lateral membrane-localized Na^+^ channels. Importantly, our work supports the hypothesis that Na^+^ channel redistribution may be a critical mechanism by which cells can rapidly respond to perturbations to support fast and robust conduction.

**Summary:** Sodium channels are organized in multiple “pools” with distinct biophysical properties and subcellular localization, but the role of these subpopulations is not well-understood. Computational modeling predicts that intercalated disk-localized sodium channels with shifted steady-state activation and inactivation voltage-dependence promote faster cardiac conduction.

## INTRODUCTION

Sodium (Na^+^) ionic current carried by voltage-gated Na^+^ channels (Nav) are primarily responsible for driving propagation of electrical activity in the heart. Na^+^ current (I_Na_) drives the rapid depolarization of myocytes that triggers the upstroke of the cardiac action potential (AP). I_Na_ dysfunction can result in pathological conditions associated with both pathological conduction and repolarization, including Brugada syndrome, long QT syndrome, atrial fibrillation, sick sinus syndrome, and heart failure^1–3^. As such, the proper expression and regulation of voltage-gated Na^+^ channels in the heart is essential for healthy cardiac function. The primary voltage-gated Na^+^ channel in cardiac myocytes is Nav1.5. A wide range of proteins have been identified to interact and regulate Nav1.5^1–3^. Different interacting proteins have been found to impact expression, trafficking, internalization, phosphorylation, and modulation of channel biophysical properties. Additionally, Nav1.5 is known to associate with potentially four different β subunits, which can also impact channel surface expression and biophysical properties, in addition to playing a role in cellular adhesion^4–7^.

Over the past two decades, multiple studies have provided evidence supporting the existence of so-called “multiple pools” of Nav1.5 in cardiac myocytes, with distinct subcellular localization, specifically referred to different subpopulations of Na^+^ channels localized at the intercalated disk (ID), i.e., at the cell-cell junctions, and at the lateral membrane^3,8–10^. Note that evidence has also suggested a third distinct subpopulation of Na^+^ channels in the transverse tubules; however, in this study, we primarily focus on ID- and lateral membrane-localized Na^+^ channels. In addition to distinct subcellular localization, these subpopulations associate with different interacting and scaffolding proteins, which supports the potential for these subpopulations to exhibit distinct biophysical properties. Indeed, macropatch measurements from Lin and colleagues showed that the ID-localized I_Na_ steady-state activation (SSA) and inactivation (SSI) voltage-dependence are shifted, compared with lateral membrane-localized I_Na_^11^. While the multiple Nav1.5 pool conceptual model is well-demonstrated across different experimental measurements, including imaging studies, protein pull-down experiments, and patch clamp^9,11–15^, the functional roles of these distinct subpopulations are not fully appreciated.

Although it is difficult to quantify the relative fraction of ID vs. lateral membrane-localized Na^+^ channels or I_Na_, measurements from Lin et al suggest that the majority of I_Na_ (relative to the entire cell) is localized at the ID. Computational modeling studies have provided strong evidence for the function role of these ID-localized Na^+^ channels in supporting and modulating electrical conduction^16–18^. At the ID, Nav1.5 has been shown to cluster with electrical and mechanical junctional proteins, primarily connexin-43 (Cx43) in the ventricles and N-cadherin, respectively, which results in the close apposition of Na^+^ channel-dense ID membranes, separated by a narrow (on the order of 5-50 nm) inter-cellular cleft space. These conditions, specifically the clustering of interacting channels adjacent to a narrow cleft space, support so-called “ephaptic coupling,” a cell-cell coupling mechanism predicted to occur in cardiac myocytes and other excitable cells^16–19^, such as neurons^20^.

In brief, this mechanism of ephaptic coupling can be described as follows: Consider electrical propagation between two coupled cells, cell 1 and cell 2, for which cell 1 is “upstream” of cell 2, i.e., propagation proceeds from cell 1 to cell 2. In cell 1, rapid inward I_Na_ at the ID membrane during the AP upstroke drives an increase in cell 1 intracellular potential; concurrently this same pre-junctional I_Na_ results in a hyperpolarization of the inter-cellular cleft space between the two cells. This cleft hyperpolarization increases the transmembrane potential across the postjunctional ID membrane in cell 2, which can promote early activation of the cell 2 post-junctional I_Na_. This cell-cell coupling occurs in parallel with direct electrical current flowing from cell 1 to cell 2 via gap junctions. The role of ephaptic coupling in cardiac electrical propagation was proposed over 50 years ago^21–24^, and subsequent computational studies have predicted that conduction depends on key properties governing ephaptic coupling, including the width of the inter-cellular cleft space and the relative fractions of I_Na_ at the ID and the lateral membrane^12,16–18,25–42^. As the clustering of Na^+^ channels at the ID occurs in conjunction with gap junctions, it is difficult to experimentally separate the relative contribution of gap junction and ephaptic coupling mechanisms, motivating computational modeling studies.

In a seminal study, Kucera and colleagues demonstrated that the impact of ephaptic coupling depended on the relative strength of gap junctional coupling^16^. Specifically, a narrow cleft space (thus promoting ephaptic coupling) enhances conduction for weaker gap junctional coupling, due to the earlier activation of post-junctional ID I_Na_ (which was later termed “self-activation”^40^). However, for stronger gap junctional, a narrow cleft can slow conduction, as the cleft hyperpolarization and post-junctional ID transmembrane potential depolarization can ultimately result in a smaller I_Na_ driving force, which in turn reduces peak I_Na_ and thus slows conduction, which was termed “self-attenuation.”^16^ Subsequent studies have found that conduction due to ephaptic coupling can further depend on additional properties, such as the relative fraction of ID-localized I_Na_ and total I_Na_ conductance (across the whole cell)^25^, and structural properties, including heterogeneous structure within different regions of the ID^26,40–42^, separation between Na^+^ channels and gap junctions on the ID membrane^40,42^, and cell size^25^.

While these computational studies have provided important insights and predictions as to how the subcellular distribution of Na^+^ channels can influence cardiac conduction via ephaptic and gap junctional coupling, all of these prior studies have assumed that the biophysical properties of Na^+^ channels localized at the ID and lateral membrane are the same. In this study, we perform a series of simulations in both single cells and tissue to predict the impact of subpopulations of Na^+^ channels with distinct biophysical properties on cardiac conduction. In tissue simulations, we consider several different subcellular distributions to identify the contributions of Na^+^ channels with distinct spatial localization, Na^+^ channels with shifted biophysical properties, and the contribution of both occurring in tandem. Collectively, our results support the conclusion that ID-localized Na^+^ channels with shifted biophysical properties enhance cardiac conduction, with enhanced sensitivity to cleft width changes (and thus greater adaptability to ID structural perturbations). Further, while simulations predict that lateral membrane-localized Na^+^ channels do contribute the total cell I_Na_, ID-localized Na^+^ channels provide the greatest contribution to the total Na^+^ charge, a property which is enhanced for weaker gap junctional coupling.

## METHODS

### Na^+^ channel biophysical properties

Simulations were performed in single cells and a one-dimensional tissue. We utilize the well-established Luo-Rudy 1991 (LR1) model^43^ to represent ionic current dynamics. All currents, except Na^+^ as described here, are unchanged from the original model. Pertinent to this study, Na^+^ channel dynamics are governed by activation, fast inactivation, and slow inactivation gates, represented by gating variables *m, h*, and *j*, respectively. These gating variables follow typical Hodgkin-Huxley type kinetics,

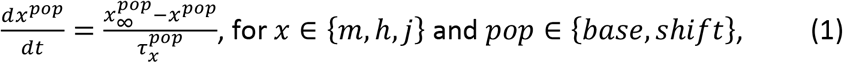

where gating variable steady-state 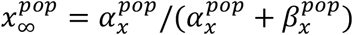 and time constant 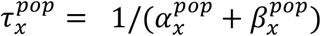 exhibit voltage-dependence through rates, 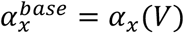 and 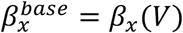, the superscript “base” here refers to the baseline model, and the functions (*α_x_*(*V*), *β_x_*(*V*)) are the nominal LR1 model voltage-dependent rate functions. The steady-steady activation (SSA) 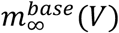 and inactivation (SSI) 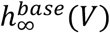 (which is equivalent to 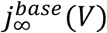) curves are shown as functions of voltage for the baseline model parameters in Figure 1A (blue).

**Figure 1.**
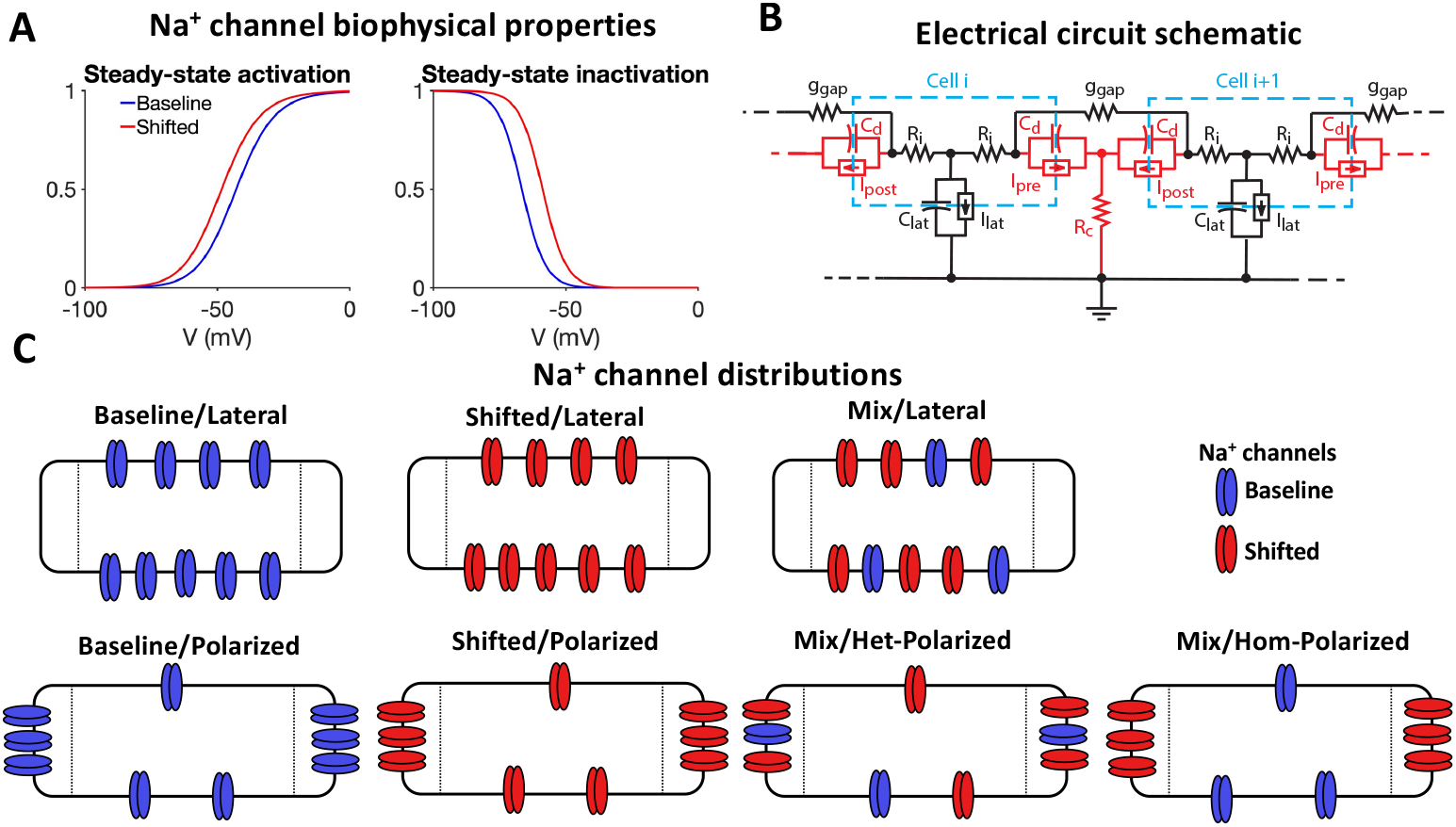
Na^+^ channel biophysical properties, electrical circuit representation, and spatial distributions. (A) The steady-state activation (left) and inactivation (right) curves, as functions of voltage V, are shown for the baseline (blue) and shifted (red) I_Na_ subpopulations. (B) Electrical circuit representation of the one-dimensional tissue model (see text for circuit element description). (C) Diagrams of the localization of baseline (blue) and shifted (red) Na^+^ channels for the 7 Na^+^ channel distributions considered. Note that the distributions in (C) correspond with f_Na_ = 2/3.

To account for a subpopulation of Na^+^ channels with different biophysical properties, we define a fraction of I_Na_ current governed by “shifted” steady-state (and time constant) voltagedependence. Specifically, we define the activation rates 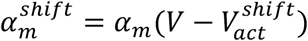 and 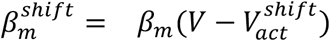, and the inactivation rates 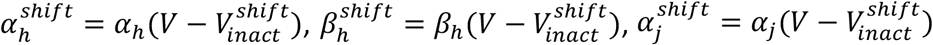, and 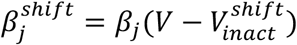. Based on measurements of the steadystate activation and inactivation curves from Lin et al^11^, we define 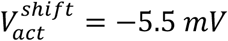 and 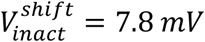. The shifted steady-steady activation 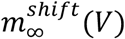 and inactivation 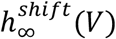 curves are shown for the shifted model parameters in Figure 1A (red). Note to account for differences in species and other experimental conditions, we incorporate the relative shift in SSA and SSI between ID and lateral membrane I_Na_, not the exact values for half-activation or - inactivation, from Lin et al^11^.

### Single cell and tissue models

In single cell simulations, we simulate the LR1 model, replacing the nominal model Na^+^ current, I_Na_ (comprised of 100% baseline I_Na_) with a sum of baseline and shifted I_Na_. We define the parameter f_Na_, which refers to the fraction of the total Na^+^ current *conductance* that has the shifted biophysical properties. That is,

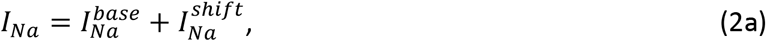

where

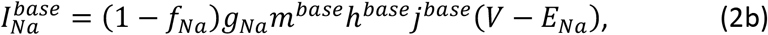

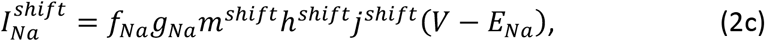

g_Na_ is the total sodium conductance and E_Na_ is the sodium reversal potential. Importantly, we highlight that, while these single cell simulations account for varying proportions of the two subpopulations with baseline and shifted I_Na_ biophysical properties, they do not account for different spatial localization of the subpopulations.

To investigate the impact of Na^+^ channel subpopulations with distinct biophysical properties and subcellular spatial locations, we simulate a tissue model of a one-dimensional chain of 50 cells, previously used by us and others^16,25,27,30,38^, which accounts for gap junction and ephaptic coupling (Figure 1B). We account for non-uniform spatial localization of Na^+^ channels by spatially discretizing each cell into three membrane “patches,” specifically the post-junctional, lateral, and pre-junctional membranes, with total membrane currents denoted as *I_post_, I_lat_*, and *I_pre_*, respectively. Thus, each cell is represented by three intracellular nodes, with intracellular resistance modeled by two resistors, *R_i_* = *ρ_i_L*/(2*πr*^2^), where *L* is the cell length of 100 μm, *r* is the cell radius of 11 μm, and intracellular resistivity *ρ_i_* = 150 Ω · cm. Capacitance of each membrane is proportional to surface area, such that the lateral membrane capacitance *C_lat_* = 2*πrLc_m_* and pre- and post-junctional intercalated disk membrane capacitance *C_d_* = *πr*^2^*c_m_*, where capacitance per unit area *c_m_* = 1 μF/cm^2^.

The dynamics of the intercellular cleft voltage (between adjacent cells) are governed by the corresponding pre- and post-junctional membrane currents *I_pre_* and *I_post_*, respectively, and the cleft radial resistance^16^ *R_c_* = *ρ_cl_*/(8*πw*), where *w* is the cleft width (varied between 5-60 nm)^4,12,30,44^, and cleft resistivity p_cl_ = 150 Ω · cm. Thus, cleft resistivity and width are inversely related. Gap junctional coupling is represented by a resistor, with conductance g_gap_, directly connecting pre- and post-junctional intracellular spaces. In this study, we vary g_gap_ from 7.67 to 767 nS^45–55^.

### Na^+^ channel distributions

Estimates based on macropatch measurements suggest that between 50-90% of Na^+^ current is localized at the ID ^11^; therefore, in this study, we consider different values for the fraction of current with shifted I_Na_ and for the fraction of current localized at the ID junctional membranes. Parameter f_Na_ defines the fraction of shifted I_Na_ and/or the fraction of junctional membrane-localized I_Na_, depending on the Na^+^ channel distribution, as described below and in Table 1.

**Table 1.** Values for the fraction of baseline and shifted I_Na_ on the lateral and junctional membranes for the 7 Na^+^ channel distributions investigated in tissue simulations, shown for a representative example of f_Na_ = 0.7.

| Distribution | Baseline $I_{Na}$ on Lateral Membrane | Shifted $I_{Na}$ on Lateral Membrane | Baseline $I_{Na}$ on Junctional Membranes | Shifted $I_{Na}$ on Junctional Membranes |
| --- | --- | --- | --- | --- |
| (i) Baseline/Lateral | 1 | 0 | 0 | 0 |
| (ii) Shifted/Lateral | 0 | 1 | 0 | 0 |
| (iii) Mix/Lateral | $0.3 (1-f_{Na})$ | $0.7 (f_{Na})$ | 0 | 0 |
| (iv) Baseline/Polarized | $0.3 (1-f_{Na})$ | 0 | $0.7 (f_{Na})$ | 0 |
| (v) Shifted/Polarized | 0 | $0.3 (1-f_{Na})$ | 0 | $0.7 (f_{Na})$ |
| (vi) Mix/Het-Polarized | $0.09 (1-f_{Na})^2$ | $0.21 [f_{Na} (1-f_{Na})]$ | $0.21 [f_{Na} (1-f_{Na})]$ | $0.49 (f_{Na})^2$ |
| (vii) Mix/Hom-Polarized | $0.3 (1-f_{Na})$ | 0 | 0 | $0.7 (f_{Na})$ |

We consider 7 distinct Na^+^ channel distributions, which collectively enable a thorough investigation of the role of different I_Na_ biophysical properties and spatial localization (Figure 1C), where we denote the fraction of the total Na^+^ current conductance of each I_Na_ subpopulation on each membrane: (i) Baseline/Lateral – all baseline I_Na_ localized on the lateral membrane; (ii) Shifted/Lateral – all shifted I_Na_ localized on the lateral membrane; (iii) Mix/Lateral – a mix of f_Na_ shifted I_Na_ and (1-f_Na_) baseline I_Na_ subpopulations, all localized on the lateral membrane; (iv) Baseline/Polarized – (1-f_Na_) baseline I_Na_ localized on the lateral membrane and f_Na_ baseline I_Na_ localized on junctional membranes; (v) Shifted/Polarized – (1-f_Na_) shifted I_Na_ localized on the lateral membrane and f_Na_ shifted I_Na_ localized on junctional membranes; (vi) Mix/Heterogeneous-Polarized – (1-f_Na_)^2^ baseline I_Na_ and f_Na_(1-f_Na_) shifted I_Na_ localized on the lateral membrane, and f_Na_(1-f_Na_) baseline I_Na_ and f_Na_^2^ localized on the junctional membrane; and (vii) Mix/Homogeneous-Polarized - (1-f_Na_) baseline I_Na_ localized on the lateral membrane and f_Na_ shifted I_Na_ localized on the junctional membrane. Note the junctional I_Na_ is evenly split between the pre- and postjunctional membranes. The fraction of the total Na^+^ current conductance of each I_Na_ subpopulation on each membrane is shown in Table 1, with the specific f_Na_ = 0.7 shown for clarity.

We highlight that the Mix/Homogeneous-Polarized distribution is consistent with a physiological representation of the I_Na_ distribution, specifically channels with shifted biophysical properties localized at the junctional membranes, while channels with baseline properties are localized on the lateral membrane. In contrast, the Baseline/Lateral distribution represents the typical model assumption, i.e., consistent with the standard monodomain model, in which all I_Na_ is represented with baseline biophysical properties and localized on the lateral membrane. The Baseline/Polarized distribution is consistent prior studies of ephaptic coupling, in which a fraction of Na^+^ channels are localized at the junctional membranes but with the same I_Na_ biophysical properties as channels localized on the lateral membrane. We also highlight the difference between the Mix/Heterogeneous-Polarized and Mix/Homogeneous-Polarized distributions. The later distribution, again representing physiological conditions, and the former distribution both have the same total fraction of I_Na_ with shifted biophysical properties and the same total fraction of I_Na_ localized at the junctional membranes. However, the Heterogeneous distribution has a mix of both I_Na_ subpopulations on both the lateral and junctional membranes, while for the Homogeneous distribution, the two I_Na_ subpopulations are localized in spatially distinct regions. Importantly, unless otherwise stated, the total I_Na_ conductance (for a given cell) is the same for all 7 Na^+^ channel distributions, with the differences only arising due to the I_Na_ subpopulations and subcellular localization.

### Simulations and numerical methods

Numerical experiments were performed by varying key model parameters in single cell and tissue simulations. In single cells, we vary the fraction of shifted I_Na_ (f_Na_) and quantify the I_Na_ peak and total Na^+^ charged carried Q_Na_. In tissue, we vary the Na^+^ channel distributions, cleft width w, gap junction conductance g_gap_, and the fraction of Na^+^ channels with shifted biophysical properties and localized at the junctional membranes (f_Na_), as described in Table 1. For each tissue simulation, we quantify the conduction velocity (CV), and in a subset of simulations, the total charged carried by I_Na_ on the lateral and junctional membranes. Unless otherwise stated, CV was measured following electrical wave initiation from resting conditions.

In single cell simulations, electrical activity initiated an applied stimulus of 1-ms duration for a cycle length of 500 ms. In tissue, a propagating wave was initiated by simulating cells 1-5. For each simulation, activation time was calculated as the first time the intracellular voltage increases above −60 mV for each cell or membrane patch (in tissue). CV was calculated by measuring the difference in activation time between the middle 50% of the one-dimensional tissue (cells 13-38). For single cells, the ordinary differential equations for gating variables and voltage are integrated using the MATLAB (The Mathworks, Inc.) ode solver *ode15s*. Tissue simulations are solved using an operator splitting method, as described in detail in our recent work^26^. In brief, gating variables for each membrane patch are integrated using forward Euler or Rush-Larson methods (for Hodgkin-Huxley type variables), which are subsequently used to update membrane ionic currents. The resulting system of ordinary differential equations and algebraic expressions governed by the electrical circuit (Figure 1B) are integrated using the backward Euler method to update intracellular and cleft voltages. For both single cells and tissue, initial conditions are set to steady-state values consistent with the resting potential of −84.5 mV.

## RESULTS

### Channels with shifted I_Na_ biophysical properties promote an earlier action potential upstroke

We first investigate the impact of heterogeneous I_Na_ populations in a single cell. Critically, in the single cell model, all ion channels are represented as being localized on the same membrane patch, i.e., there is no I_Na_ spatial distribution. We first consider the case of a mixed 50:50 I_Na_ distribution (f_Na_ = 0.5), equally split between baseline and shifted I_Na_ (magenta [voltage traces], red/blue [I_Na_ traces]) and compare with the nominal homogeneous model, with only baseline I_Na_ (black). At the level of the single cell AP, we find almost no difference between the cell with the mixed I_Na_ subpopulations and only baseline I_Na_, except for small changes in the peak voltage (Figure 2A). However, closer examination reveals that the presence of the shifted I_Na_ results in an earlier AP upstroke (magenta), compared with the only baseline I_Na_ model (black). We find that the early AP upstroke is due to the subpopulation of shifted I_Na_ activating earlier (red), which in turn activates the subpopulation of baseline I_Na_ earlier (blue), compared with the only baseline I_Na_ model (black). Additionally, in the cell with both subpopulations, the peak of the shifted I_Na_ is greater than the peak of the baseline I_Na_. That is, even though the conductances of the two subpopulations are equal, the magnitude of the shifted I_Na_ subpopulation is larger compared with the baseline I_Na_.

**Figure 2.**
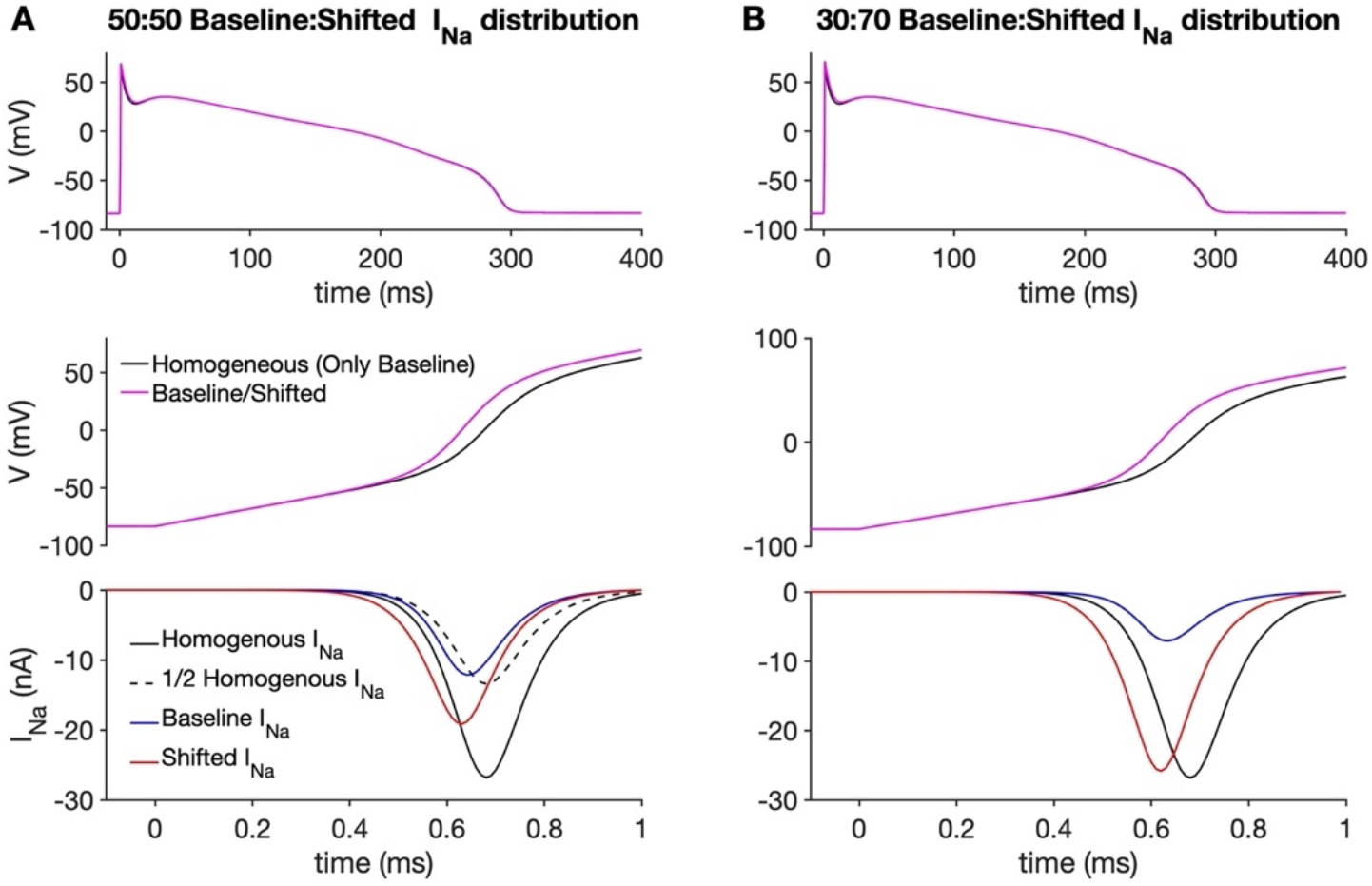
Addition of a subpopulation of I_Na_ with shifted biophysical properties results in earlier AP upstroke and larger I_Na_ current in a single cell. Transmembrane voltage (V) (row 1, 2; magenta) and I_Na_ (row 3; blue/red) are shown as functions of time for a single cell with a (A) 50:50 and (B) 30:70 distribution of baseline:shifted I_Na_ subpopulations. The corresponding voltage and I_Na_ curves are shown for the homogeneous (only baseline I_Na_) single cell in black. In (A), I_Na_ scaled by a factor of 1/2 is shown for comparison with I_Na_ from the baseline/shifted single cell (dashed black line).

In a single cell with 70% shifted I_Na_ (f_Na_ = 0.7), we find that the effects described above are magnified (Figure 2B). The AP upstroke is earlier, compared with the 50-50% distribution, and the shifted I_Na_ subpopulation is greatly enhanced and is almost equal in magnitude (red) to the only baseline I_Na_ (black).

We next quantified how the I_Na_ peak and total Na^+^ charge (Q_Na_) for the two I_Na_ subpopulations varies, as the fraction of the shifted I_Na_ subpopulation changes in a single cell (Figure 3). We first measure the ratio of the shifted I_Na_ peak to the baseline I_Na_ peak as a function of f_Na_ (Figure 3A). As f_Na_ increases, the shifted I_Na_ peak ratio also increases, as expected. However, we can compare this increase with expectations based solely on the change in conductance between the two subpopulations. Based on conductance changes alone, we would expect the ratio between the two peaks to scale with f_Na_/(1-f_Na_) (dashed line). Critically, the actual simulated ratio (black) is always greater than this expectation, meaning that the shifted I_Na_ peak is always proportionally larger than the shifted I_Na_ conductance.

**Figure 3.**
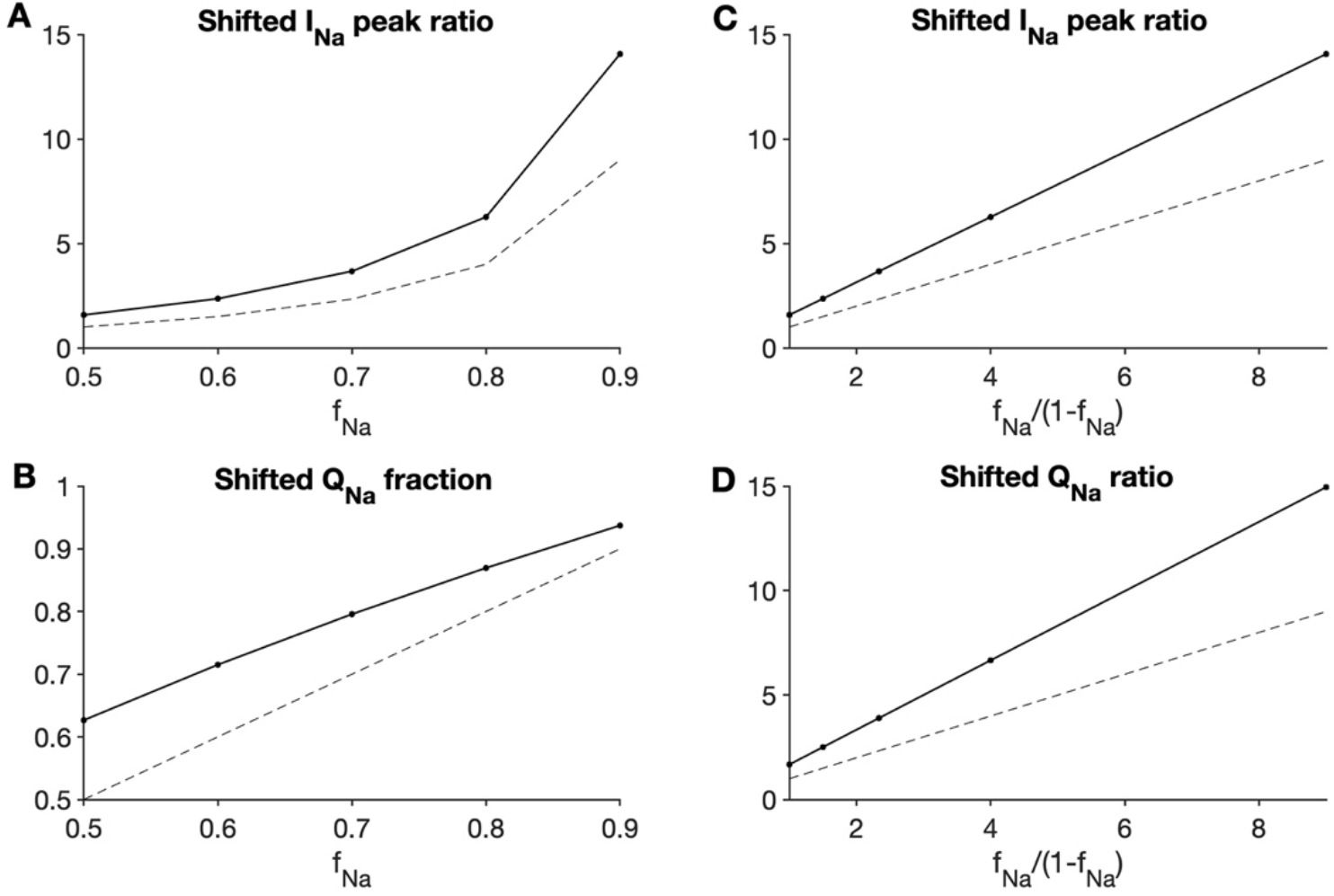
Na^+^ current with shifted biophysical properties is proportionally larger and contributes more charge than its conductance fraction in single cells. (A) The ratio of the shifted I_Na_ peak-to-baseline I_Na_ peak and (B) the fraction of charge carried by the shifted I_Na_ subpopulation Q_Na_ are shown as a function of the shifted I_Na_ conductance fraction (f_Na_). (C) The shifted I_Na_ ratio and (D) shifted Q_Na_ ratio are shown as a function of the shifted I_Na_-to-baseline I_Na_ conductance fraction ratio (f_Na_/[1-f_Na_]).

We also quantified the total charge carried by the two I_Na_ subpopulations and measured the fraction of the total Na^+^ charge (Q_Na_) carried by the shifted I_Na_ subpopulation (Figure 3B). Based on conductance alone, we would expect this fraction to scale with f_Na_ (dashed line). However, we find that the shifted Q_Na_ (black) is always greater than this expectation as well. That is, the shifted I_Na_ subpopulation carries a proportion of the total Na^+^ charge greater than its respective conductance. Plots of the I_Na_ and Q_Na_ ratios against f_Na_/(1-f_Na_) similarly show that the shifted I_Na_ subpopulation proportionally contributes more to the total Na^+^ current, compared with the baseline I_Na_, than expected based on conductance (Figure 3C, D).

### Shifted I_Na_ subpopulation enhances conduction in cardiac tissue

The above results demonstrate that in a single cell, channels with shifted I_Na_ biophysical properties contribute to an earlier AP upstroke. We next systematically investigate to what extent conduction in a one-dimensional tissue depends on the presence of Na^+^ channels with shifted I_Na_ and their localization at the ID. Further, we investigate how these properties also depend on ID structure, specifically the intercellular cleft width and the strength of gap junctional coupling.

As described in the Methods, we consider 7 distinct Na^+^ channel distributions, chosen to elucidate the role of both the presence of channels with shifted I_Na_ biophysical properties and their localization at the ID. In Figure 4, we plot CV as a function of the cleft width for different values of the gap junction conductance *g_gap_*. As expected, for all Na^+^ channel distributions, CV increases with increasing *g_gap_*. For the three distributions with only Lateral Na^+^ channels (Figure 4A-C), CV does not depend on cleft width, also as expected since these distributions do not engage ephaptic coupling. Of these three Lateral distributions, conduction is fastest in the tissue with only shifted I_Na_ (Figure 4B), slightly faster than in tissue with a 30:70 Baseline:Shifted Mixed I_Na_ distribution (Figure 4C), with the only baseline I_Na_ slowest (Figure 4A).

**Figure 4.**
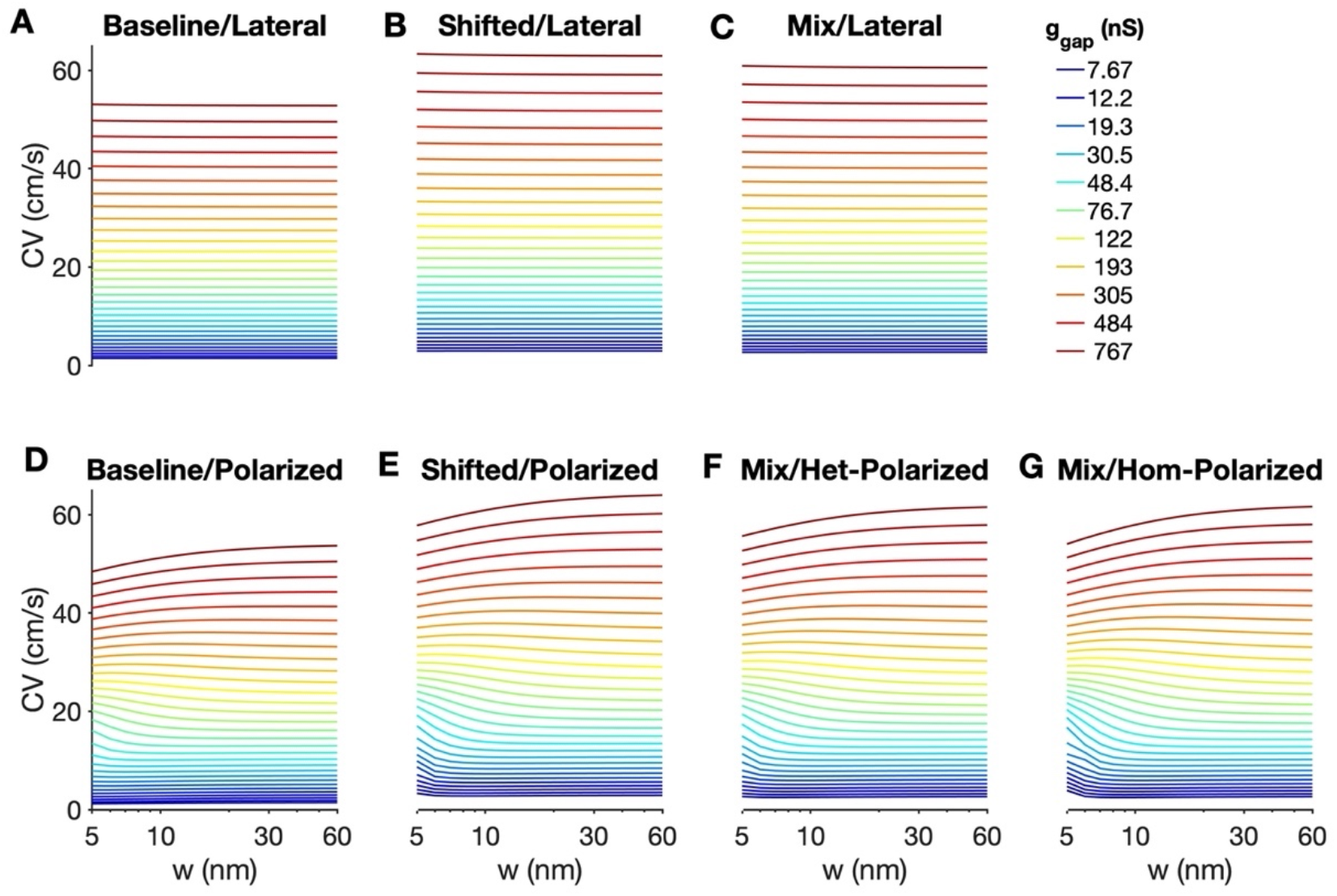
Conduction velocity depends on Na^+^ channel distribution, cleft width, and gap junctional coupling. Conduction velocity (CV) is shown as a function of cleft width (w) for different Na^+^ channel distributions and gap junction conductance g_gap_. For Polarized distributions, narrow cleft slows conduction for high g_gap_ and enhanced conduction for low g_gap_. The physiological distribution (Mix/Homogeneous-Polarized) exhibits the greatest sensitivity to cleft width.

The presence of the shifted I_Na_ subpopulation in the Polarized distributions similarly results in faster conduction. In contrast with the Lateral distributions, the four Polarized distributions (Figure 4D-G) exhibit a clear dependence on the cleft width. Consistent with prior studies^16,17,25,27,33^, for all Polarized distributions, narrowing the cleft tends to slow conduction for stronger gap junctional coupling, while narrowing the cleft tends to enhance conduction for weaker gap junctional coupling. Further, we find that the tissue with Shifted/Polarized and Mixed/Polarized I_Na_ distributions exhibit greater sensitivity to the cleft width, compared with the Baseline/Polarized distribution.

In Figure 5, we plot the same CV measurements, as a function of g_gap_ for different values of cleft width w. The Lateral I_Na_ distributions similarly exhibit no dependence on cleft width (Figure 5A-C). From this presentation, it is also more clear to what extent CV varies for the four Polarized distributions, as a function of cleft width and g_gap_ (Figure 5D-G). As noted above, cleft narrowing results in condution slowing for high g_gap_ and conduction enhancing for low g_gap_. Importantly, we find that the Mixed/Homogeneous-Polarized distribution depends on cleft width for the widest range of g_gap_ values and demonstrates the greatest sensitivity to cleft width (for a given g_gap_).

**Figure 5.**
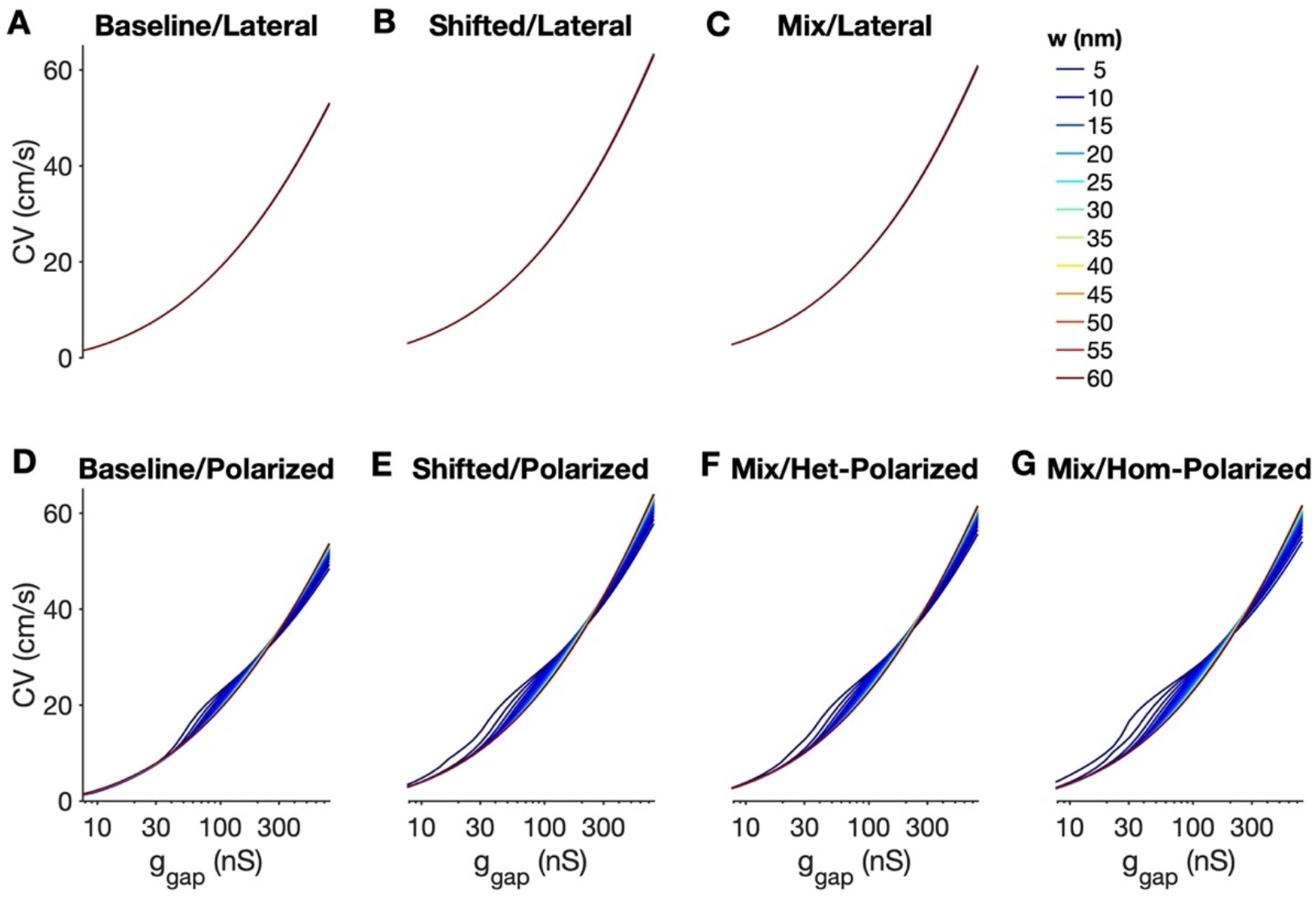
Conduction velocity depends on Na^+^ channel distribution, gap junctional coupling, and cleft width. Conduction velocity (CV) is shown as a function of gap junction conductance g_gap_ for different Na^+^ channel distributions and cleft width w. CV slows for decreasing g_gap_, with sensitivity to cleft width greater for polarized distributions with the shifted and mix I_Na_ subpopulations.

In Figure 6, we plot CV as a function of g_gap_ and directly compare a subset of the Na^+^ channel distributions. For all cleft widths and most g_gap_ values (except the extreme high and low g_gap_), conduction is fastest for the Mixed/Homogeneous-Polarized (blue), followed by the Mixed/Heterogeneous-Polarized (magenta), Mixed/Lateral (black), Baseline/Polarized (red), and Baseline/Lateral (green). We also show ratios of the CV values for several combinations of Na^+^ channel distributions, to identify to what extent Na^+^ channel composition vs. localization regulate conduction. Comparing the Polarized vs Lateral Distribution with Baseline I_Na_, the Polarized distribution has faster CV, i.e., ratio > 1, for a range of g_gap_ values between about 50-500 nS, but slower CV for larger or smaller g_gap_ outside this range (Figure 6B, solid black). Comparing the Polarized vs Lateral Distribution for the Mixed I_Na_ distributions, the Polarized distributions is similarly faster for a g_gap_ range and slower for outside the range (Figure 6B, dashed black). However, the g_gap_ range for enhanced CV is widened to about 20-500 nS. That is, polarizing the mixed I_Na_ subpopulation enhances conduction for a wider range of g_gap_.

**Figure 6.**
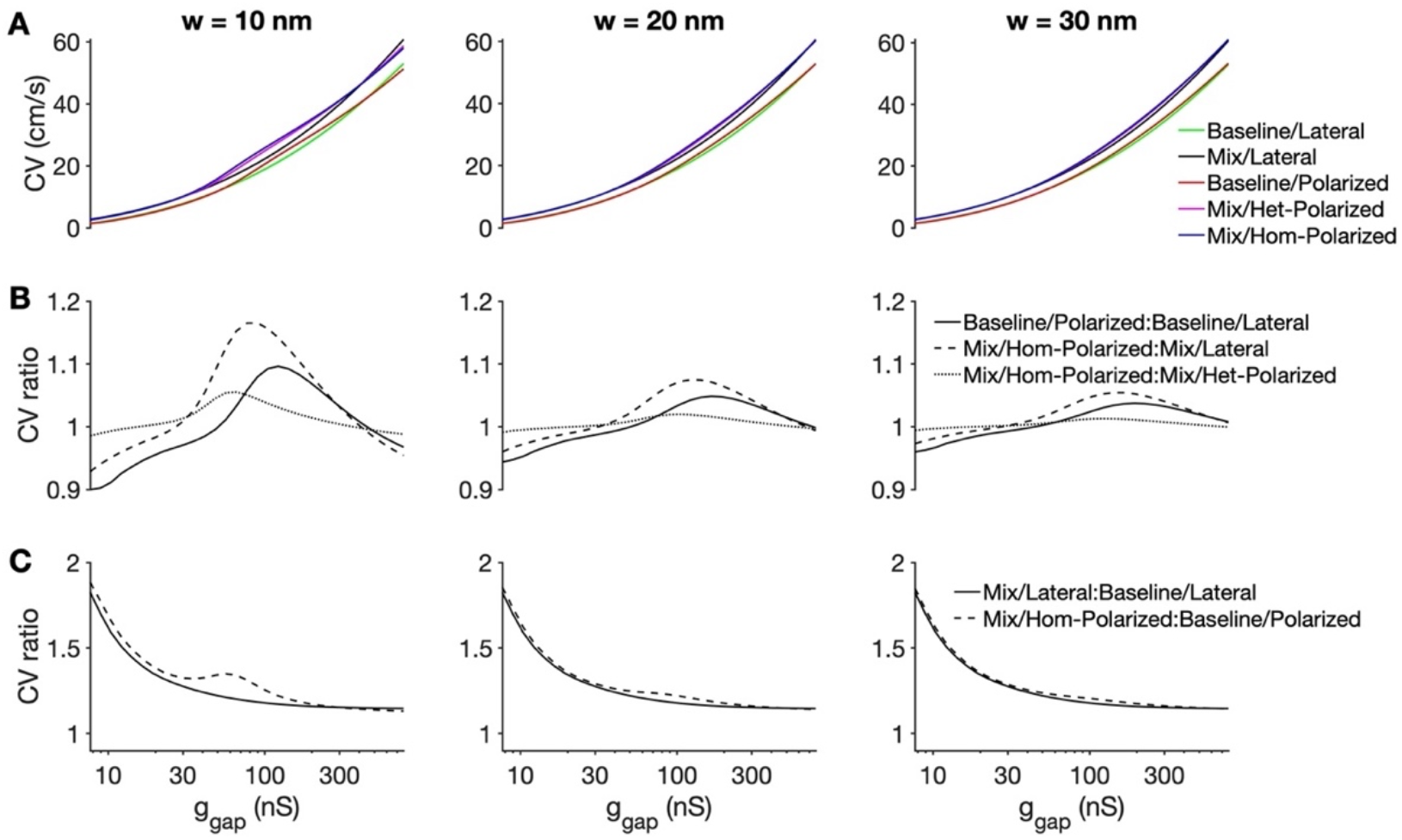
The physiological (Mix/Homogeneous-Polarized) distribution exhibits enhanced conduction. (A) Conduction velocity (CV) is shown as a function of gap junction conductance g_gap_ for cleft widths of 10 (left), 20 (middle), and 30 (right) nm and different Na^+^ channel distributions. (B, C) The ratio of CV for different Na^+^ channel distribution combinations are shown. Parameters: f_Na_ = 0.7.

We can also directly compare the Mixed/Homogeneous-Polarized vs Mixed/Heterogeneous-Polarized distributions, which have the same fraction of baseline vs shifted I_Na_ current and the same fraction of I_Na_ polarized at the ID, with the only difference being whether or not the I_Na_ subpopulations are spatially distinct. We find that for nearly all g_gap_ values, this CV ratio > 1, i.e., polarization specifically of the shifted I_Na_ subpopulation enhances conduction (Figure 6B, dotted line). Comparing the Mixed/Lateral vs Baseline/Lateral distribution, we find that the CV ratio > 1 for all g_gap_ values, and the ratio is increases as g_gap_ is decreased (Figure 6C, solid line). That is, with only Lateral Na^+^ channels, the Mixed I_Na_ subpopulations always enhances conduction, relative to Baseline I_Na_, and to a greater extent for weak gap junctional coupling. Interestingly, when we make the comparable comparison in Polarized tissue (Mixed/Homogeneous-Polarized vs. Baseline/Polarized), we observe the same general trend, with the addition of an additive “bump” between g_gap_ of 30 and 200 nS (Figure 6C, dashed line). Across all distributions and g_gap_, the magnitude of these effects generally decreases as cleft width w increases.

Collectively, these results support the following general conclusions: (i) the inclusion of a subpopulation with shifted I_Na_ biophysical properties speeds up conduction, with CV enhancement increasing as gap junctional coupling weakens; (ii) polarization of any Na^+^ channels enhances conduction over a wide range of gap junctional coupling levels; (iii) polarization of the shifted I_Na_ subpopulation results in an additional conduction enhancement (over a wider g_gap_ range); and (iv) the influence of Na^+^ channel polarization on conduction depends on the cleft width, while the role of shifted I_Na_ subpopulation predominantly does not depend on cleft width (except in the 30-100 nS g_gap_ range).

In Supporting Figures S1 and S2, we plot these same CV ratios for different combinations of Na^+^ channel distributions for different values of f_Na_. For nearly all cases, the magnitude of the relationships described above are larger as f_Na_ is increased. That is, the influence of the shifted I_Na_ subpopulation and the polarization of specific Na^+^ channel subpopulations is magnified as the subpopulation fraction is increased.

In Figure 7, we show contour maps of CV for the Baseline/Polarized, Mix/Homogeneous-Polarized, and Shifted/Polarized distributions as functions of cleft width and f_Na_ for different values of gap junction conductance g_gap_. Importantly, these maps illustrate the range of CV changes due solely to tissue structural changes (i.e., cleft width variation) and Na^+^ channel subpopulation redistribution (i.e., varying f_Na_) for a fixed degree of gap junctional coupling. Several key points are apparent from these plots: For all conditions, CV is larger for the Mix/Homogeneous-Polarized distribution, relative to Baseline/Polarized, but slower than the Shifted/Polarized distribution. Further, CV depends on both cleft width and f_Na_; however, this relationship depends on gap junctional coupling. For low g_gap_, CV predominantly depends on cleft width, while for high g_gap_, CV predominantly depends on f_Na_, with moderate g_gap_ exhibiting comparable dependence on both. Thus, for weak gap junctional coupling, CV is most effectively modulated by changes in cleft width, while for strong gap junctional coupling, CV is most effectively modulated by Na^+^ channel redistribution. Additionally, the number of contour regions (with 1-cm/s spacing) illustrates that CV for the Mix/Homogenous-Polarized distribution exhibits the greatest variability, compared with either the Baseline/Polarized or Shifted/Polarized distributions. That is, the physiological distribution, through cleft width changes and Na^+^ channel redistribution, can result in a wider range of CV values, for fixed gap junctional coupling. Further, this variability in CV is wider for moderate g_gap_, i.e., CV can be modulated to the greatest degree for moderate levels of gap junctional coupling.

**Figure 7.**
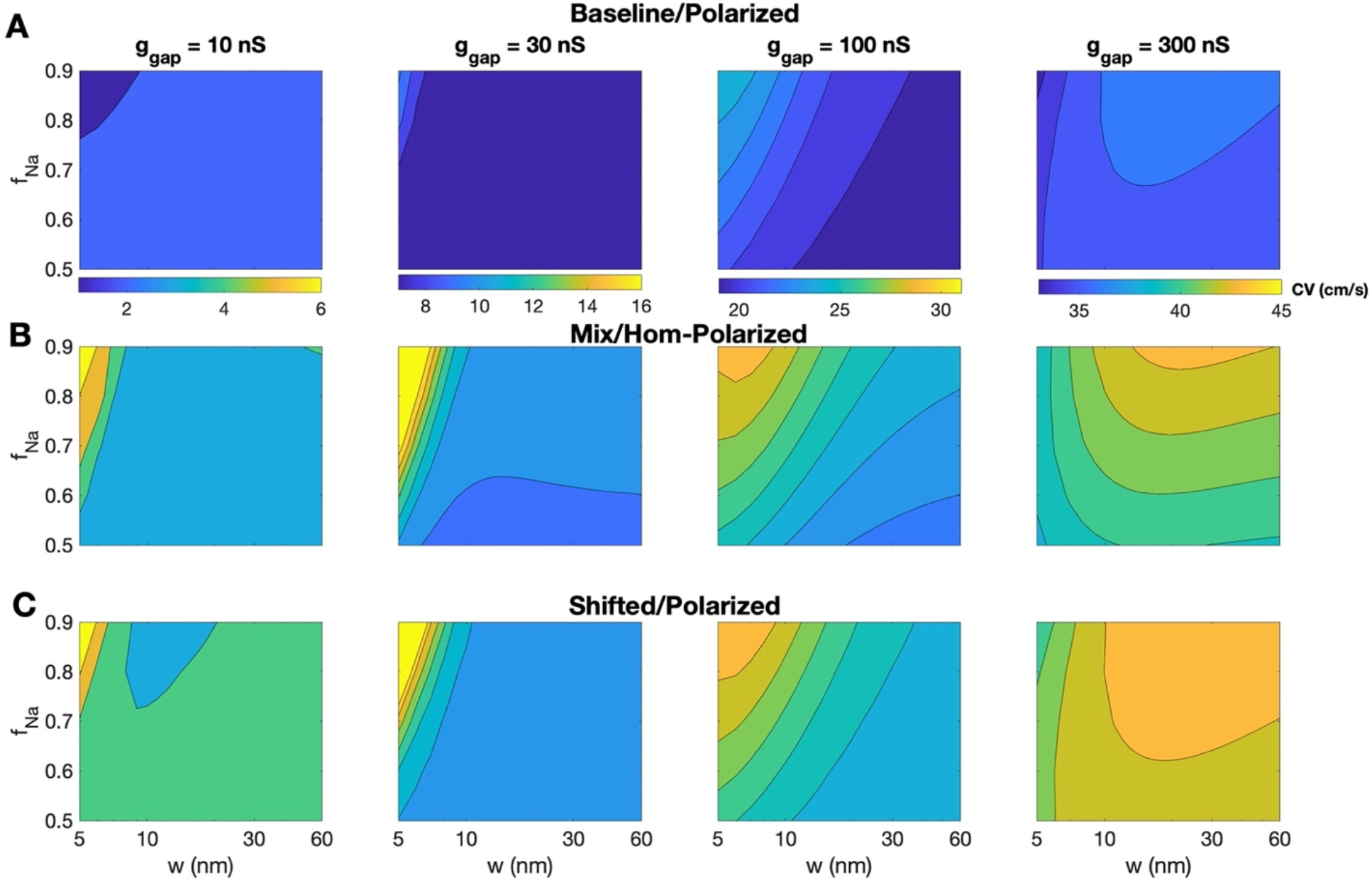
The physiological (Mix/Homogeneous-Polarized) distribution exhibits sensitivity to cleft widths and Na^+^ channel redistribution. Contour maps of the conduction velocity (CV) are shown for the (A) Baseline/Polarized, (B) Mix/Homogeneous-Polarized, and (C) Shifted/Polarized distributions as functions of cleft width w and f_Na_, with 1-cm/s spacing.

### Shifted I_Na_ subpopulation promote larger faster cell-cell transmission and post-junctional I_Na_

We next sought to understand the mechanisms underlying the changes in conduction due to Na^+^ channel biophysical properties and distribution. In Figure 8A (top), we plot the activation time for the Baseline/Polarized (black) and Mix/Homogeneous-Polarized (magenta) distributions as a function of the cell number, for which the time of post-junctional, lateral, and pre-junctional membrane activation is denoted for each cell. Both simulations have moderate gap junctional coupling (g_gap_ = 100 nS) and I_Na_ polarization (f_Na_ = 0.7). Consistent with the faster CV measurements above, activation time is earlier for the Mix/Homogeneous-Polarized distribution. For direct comparison, we also shifted the curves, such that the two align at the activation time of either the pre-junctional membrane of cell 24 (Figure 7A, middle) or post-junctional membrane of cell 25 (bottom). From these two plots, we can identify that the primary difference in activation time is due to the delay in time between the cell 24 pre-junctional membrane and the cell 25 post-junctional membrane, i.e., the time for the electrical wave to propagate across the cell-cell junction. In contrast, the delay between the post-junctional and pre-junctional membrane of cell 25 is near identical for the two distributions, i.e., there is no difference in the time for the electrical wave to propagate across the intracellular space.

**Figure 8.**
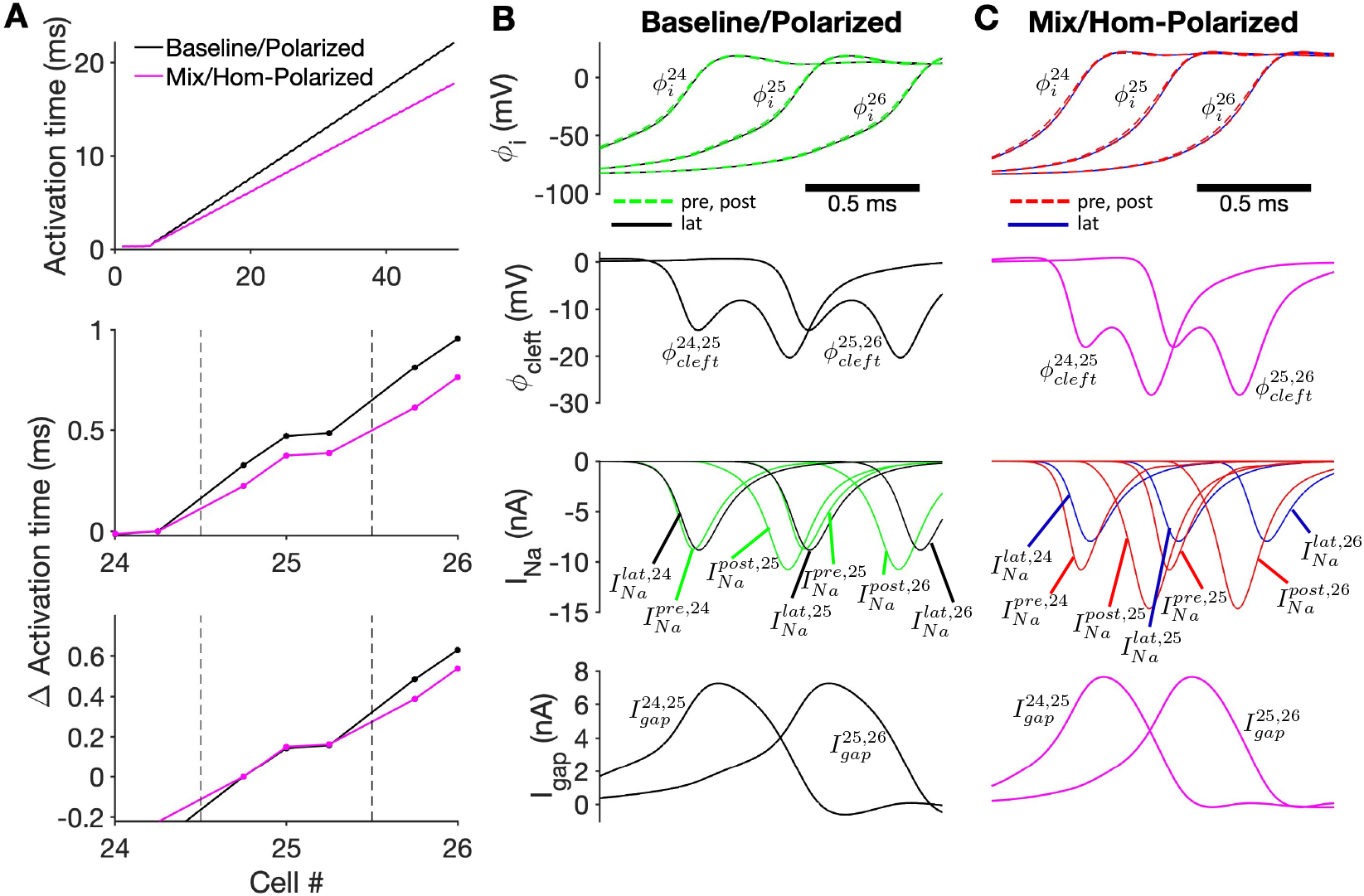
Enhanced post-junctional I_Na_ with shifted biophysical properties promotes faster cell-cell transmission and conduction. (A) (top) Activation time for post-junctional, lateral, and pre-junctional membranes for each cell for the Baseline/Polarized (black) and Mix/Homogeneous-Polarized (magenta) distributions. The change in activation time is shown, adjusted to (middle) cell 24 pre-junctional membrane and (bottom) cell 25 post-junctional membrane activation times. (B, C) Intracellular potential φ, (row 1); cleft potential φ*cieft* (row 2); pre- and post-junctional I_Na_ (B, green; C, red) and lateral I_Na_ (B, black; C, blue) (row 3); and gap junction current (row 4) are shown as functions of time for (B) Baseline/Polarized and (C) Mix/Homogeneous-Polarized distributions. Parameters: g_gap_ = 100 nS, f_Na_ = 0.7, w = 10 nm.

We next investigate the voltage and I_Na_ changes associated with these two cases (Figure 8B, C). We observe a slower AP upstroke in the Baseline/Polarized distribution, compared with the Mix/Homogeneous-Polarized distribution (Figure 8B, C, row 1), consistent with the single cell simulations in Figure 2. In both distributions, the cleft space between the cells is hyperpolarized, consistent with prior studies of ephaptic coupling mechanisms^16,25,26,40,42^ (Figure 8B, C, row 2). However, we find a larger magnitude hyperpolarization in the Mix/Homogeneous-Polarized distributions (magenta). Further, while there are small differences in time course and magnitude, the gap junctional current is comparable in both distributions, highlighting that the differences in conduction velocity are directly attributable to the differences in Na^+^ channel distribution.

However, examination of the I_Na_ reveals critical differences between the two distributions, specifically considering the cell 24 lateral and pre-junctional I_Na_; cell 25 post-junctional, lateral, and pre-junctional I_Na_; and cell 26 post-junctional and lateral I_Na_. For the Baseline/Polarized distribution, for a given cell (here, cell 25) we find that the post-junctional I_Na_ (green) activates earliest, as expected. However, the lateral (black) and pre-junctional (green) I_Na_ activate at almost the same time, with the pre-junctional I_Na_ peak slightly earlier. Additionally, the post-junctional peak I_Na_ is largest in magnitude, while the lateral and pre-junctional peak I_Na_ are almost identical magnitude. For the Mix/Homogeneous-Polarized distribution, again for a given cell (considering cell 25), the post-junctional I_Na_ (red) similarly activates earliest and has the largest peak. However, the pre-junctional I_Na_ (red) activates next and has the second largest peak, while the lateral I_Na_ (blue) activates last and has the smallest peak. That is, due to the shifted I_Na_ biophysical properties of the junctional I_Na_ current, the pre-junctional I_Na_ activates before the lateral membrane, despite being further ‘downstream’ in the cell. In Supporting Figure S3, we quantify the delay between the timing of the I_Na_ peak for the post-junctional, lateral, and pre-junctional I_Na_ for cell 25 and the AP activation of the cell 24 post-junctional membrane, for different values of f_Na_ and cleft width. We find that the post-junctional I_Na_ peak is consistently earliest and increasingly so for larger f_Na_ and smaller cleft width. For nearly all cases and both distributions, the pre-junctional I_Na_ peak is earlier than the lateral membrane. However, the difference in peak timing is greatest for the Mix/Homogeneous-Polarized distribution, compared with the Baseline/Polarized distribution, and for larger f_Na_ and smaller cleft width.

For the value of f_Na_ = 0.7 (Figure 8), the total I_Na_ conductance distribution is thus 35%, 30%, and 35% for the post-junctional, lateral, and pre-junctional membranes, respectively. However, for both Na^+^ channel distributions considered in Figure 8, the post-junctional I_Na_ is disproportionally larger, compared to this expectation based on conductance. Further, for the Mix/Homogeneous-Polarized distribution, the lateral I_Na_ is disproportionally smaller. That is, the pre-junctional I_Na_ contributes more to the total I_Na_, while the lateral I_Na_ contributes less, than simply the expected proportion based on the conductance distribution.

We quantify these trends by plotting the fraction of total charge carried by I_Na_ on each membrane (Q_Na_) as a function of f_Na_ and for different cleft widths (Figure 9). Based on conductance distribution, the post- and pre-junctional membranes would be expected to scale with f_Na_/2, while the lateral membrane would be expected to scale with 1-f_Na_, shown as dashed black lines. As expected, both post- and pre-junctional Q_Na_ increase with f_Na_, while lateral Q_Na_ decreases with f_Na_. We find that for both distributions, post-junctional Q_Na_ is larger than expectation based on the conductance distribution (Figure 9A), with Q_Na_ for the Mix/Homogeneous-Polarized distribution consistently larger than the Baseline/Polarized distribution. Both of these trends are greatest for narrow cleft width.

**Figure 9.**
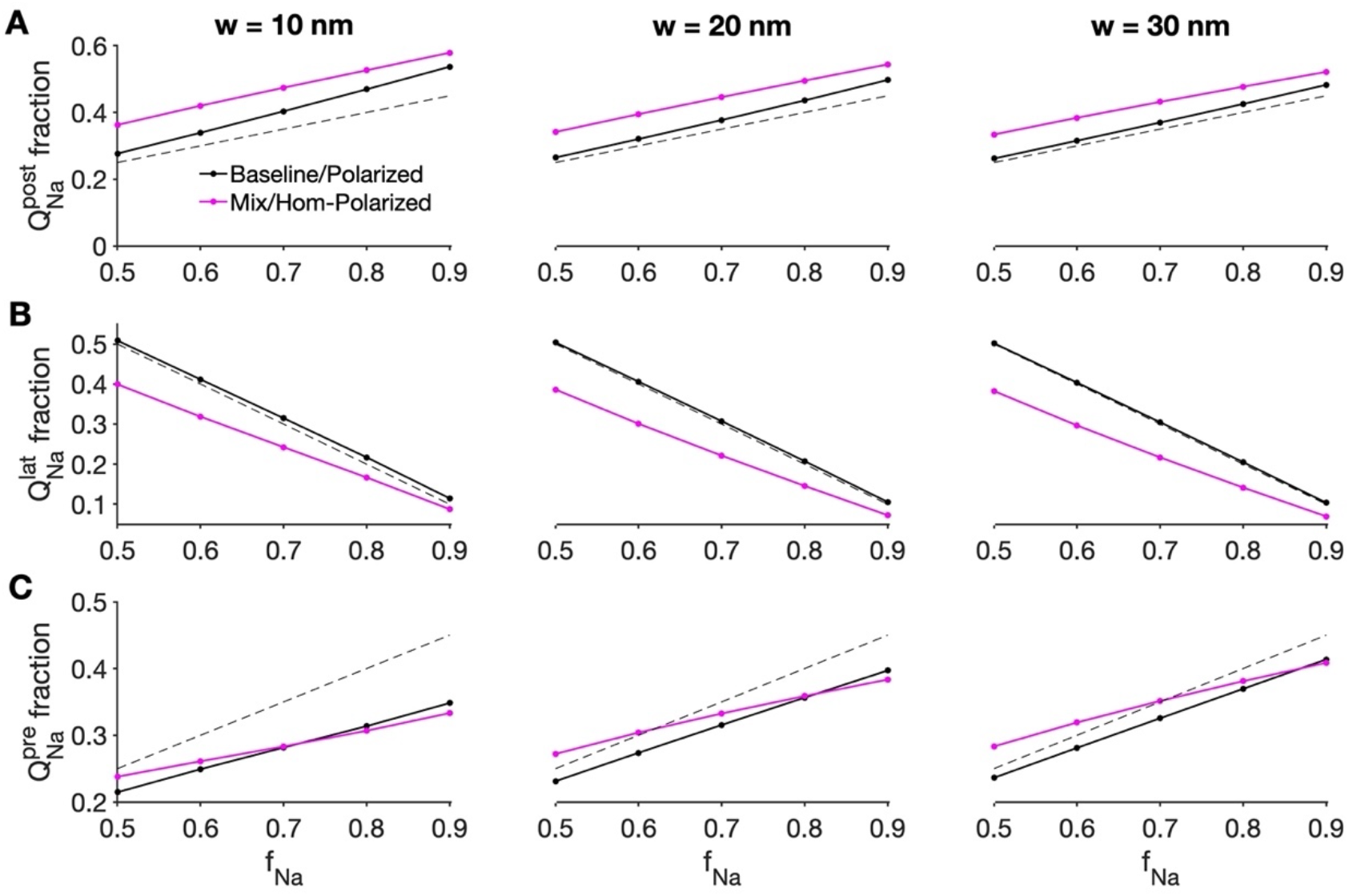
Post-junctional I_Na_ contributes proportionally more charge than the conductance fraction. The fraction of Na^+^ charge Q_Na_ carried by (A) post-junctional, (B) lateral, and (C) pre-junctional I_Na_ is shown as a function of f_Na_, for cleft widths of 10 (left), 20 (middle), and 30 (right) nm and different Na^+^ channel distributions (Baseline/Polarized, black; Mix/Homogeneous-Polarized, magenta). The dashed black line corresponds with the conductance fraction on each membrane (f_Na_/2 on post- and pre-junctional; 1-f_Na_ on lateral membrane). Parameters: g_gap_ = 100 nS.

Interestingly, the lateral Q_Na_ for the Baseline/Polarized distribution is nearly identical to the conductance expectation (dashed line), i.e., the lateral I_Na_ charge is proportional to its conductance (Figure 8B). In contrast, lateral Q_Na_ for the Mix/Homogeneous-Polarized distribution is less than the conductance distribution. Due the opposing trends for the post-junctional and lateral Q_Na_, the pre-junctional Q_Na_ fraction trends are inconsistent. For smaller f_Na_, the Mix/Homogeneous-Polarized distribution pre-junctional Q_Na_ fraction is larger than the Baseline/Polarized distribution, and to a greater extent for wider cleft width.

Thus, in summary, for both distributions, the post-junctional I_Na_ is consistently contributing the largest fraction of the total I_Na_ charge, with a greater contribution for the Mix/Homogeneous-Polarized distribution. As a consequence, the lateral membrane I_Na_ contribute proportionally contribute less to the total I_Na_, more so for tissues with the Mixed subpopulation.

In Figure 10, we plot the Q_Na_ fractions as functions of g_gap_ for different values of f_Na_ and cleft width for the Mix/Homogeneous-Polarized distribution. For all cleft width and g_gap_, post- and prejunctional Q_Na_ increase with increasing f_Na_, while lateral Q_Na_ decreases, as expected. However, Q_Na_ fractions exhibit an interesting dependence on g_gap_. For all cases, the lateral Q_Na_ decreases as g_gap_ decreases (Figure 10B). However, post-junctional Q_Na_ exhibits a biphasic dependence on g_gap_, with a positive ‘bump’ between around 30 and 300 nS (Figure 10A), while pre-junctional Q_Na_ exhibits a similar negative ‘bump’ in the same g_gap_ range (Figure 10C). Thus, collectively these results demonstrate that as gap junctional coupling weakens, lateral Q_Na_ decreases, such that the two junctional membranes constitute a larger Q_Na_ contribution. Further, the post-junctional Q_Na_ is always larger than the pre-junctional Q_Na_. However, this disparity is greatest for g_gap_ values around 30-300 nS, with pre-junctional Q_Na_ disproportionally largest, while for extreme cases of either high or low gap junctional coupling, post- and pre-junctional Q_Na_ are much similar in magnitude. The magnitude of these trends is generally smaller for wider cleft width. In Supporting Figure S4, we observed similar trends for the Baseline/Polarized distribution, with the lateral Q_Na_ generally contributing a larger proportion, consistent with the result in Figure 10.

**Figure 10.**
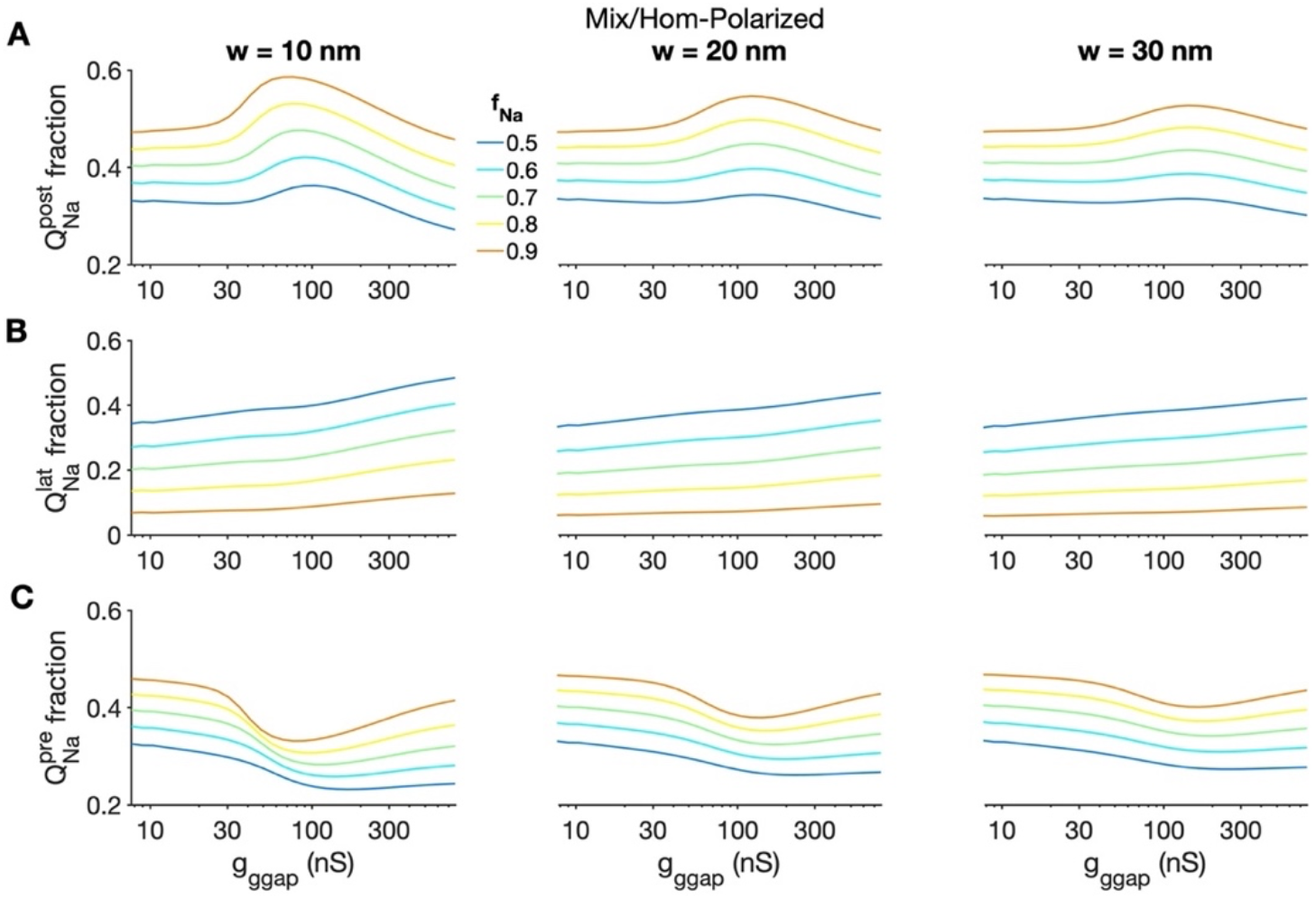
Post-junctional I_Na_ contribution is enhanced for moderate gap junction coupling. The fraction of Na^+^ charge Q_Na_ carried by (A) post-junctional, (B) lateral, and (C) pre-junctional I_Na_ is shown as a function of gap junction conductance g_gap_ for cleft widths of 10 (left), 20 (middle), and 30 (right) nm and different values of f_Na_ for the Mix/Homogeneous-Polarized distribution.

To further demonstrate the functional role of the ID-localized I_Na_, we measure CV for membrane-specific block in the Mix/Homogeneous-Polarized distribution for f_Na_ = 0.7 (Figure 11). Specifically, we reduce the I_Na_ conductance on either the ID (red) or lateral (blue) membranes, while maintaining the same conductance on the other membrane. We consider the full range from the nominal Mix/Homogeneous-Polarized distribution with 70% ID-localized shifted Na^+^ channels and 30% lateral membrane-localized baseline Na^+^ channels and decrease one of these subpopulations individually until only the other subpopulation is present. We note that this results in a decrease in overall cell I_Na_ conductance. We find that reducing the conductance of either the ID- or lateral membrane-localized I_Na_ slows conduction. However, across all conditions (cleft width and gap junction conductance), we find that reducing ID-localized I_Na_ results in greater conduction slowing, with generally a greater disparity for larger g_gap_. In Supporting Figure S5, we find the same trends for f_Na_ = 0.5.

**Figure 11.**
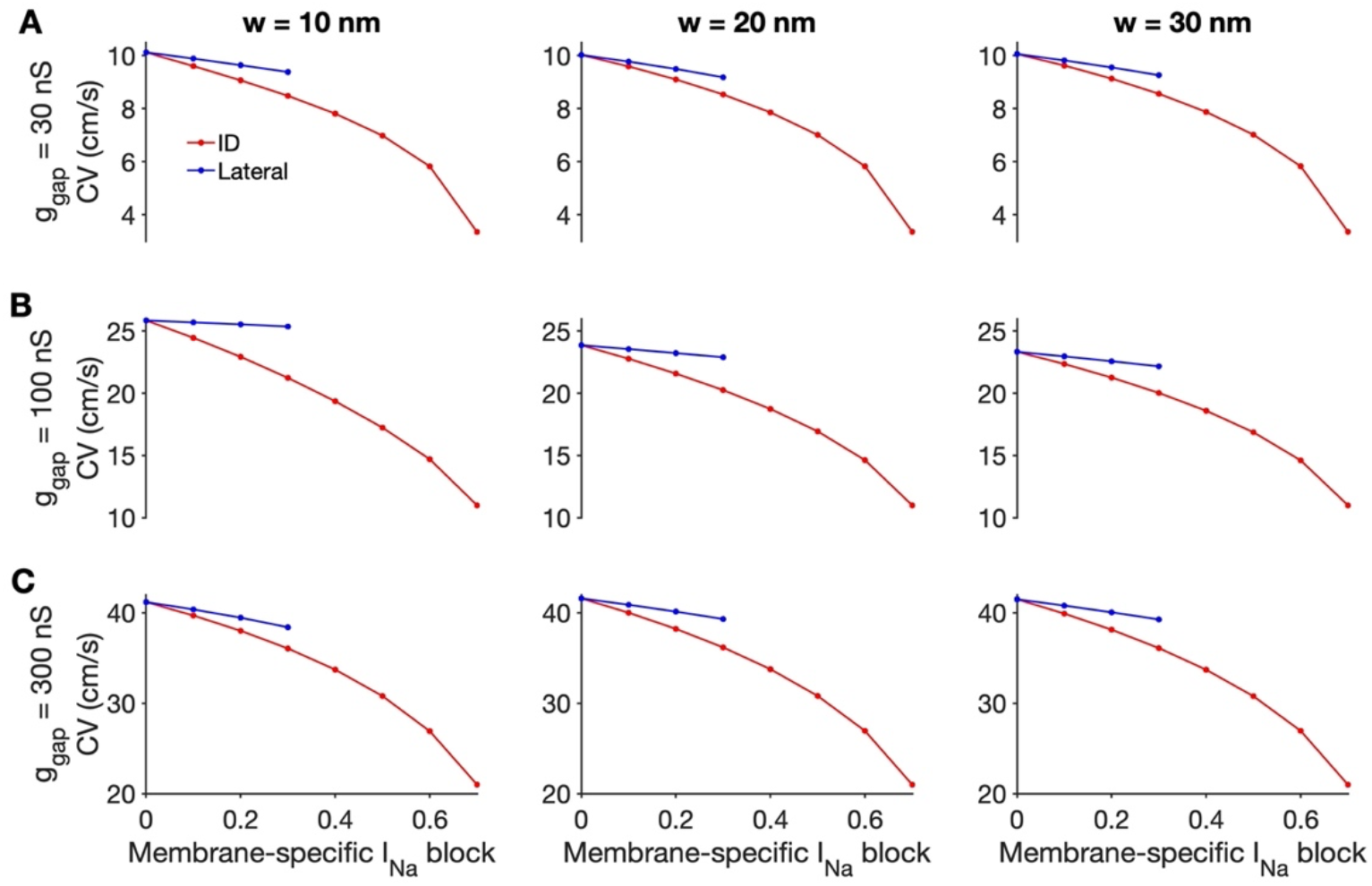
Membrane-specific I_Na_ block in the Mix/Homogeneous-Polarized distribution. (A-C) Conduction velocity (CV) is shown as a function of the I_Na_ conductance block of the ID (red) or lateral (blue) membrane, for different values of cleft width w and gap junction conductance g_gap_. Parameters: f_Na_ = 0.7.

All of the previous simulations considered conduction from rest. We next consider how different Na^+^ channel distributions impact conduction for faster pacing rates, i.e., shorter basic cycle length (BCLs). For the Baseline/Polarized and Mix/Homogeneous-Polarized distributions, we paced the tissues at different BCL values until steady-state or a steady-state alternating pattern was reached and measured CV for different cleft widths and g_gap_ (Figure 12). Consistent with the above results, across all BCLs, conduction is faster for the Mix/Homogeneous-Polarized distribution (magenta), compared with the Baseline/Polarized distribution (black). Additionally, for all cleft widths and g_gap_, tissue with the Mix/Homogeneous-Polarized distribution could be paced at shorter BCLs, before loss of 1:1 capture or conduction block. Additionally, for faster pacing rates, the tissues exhibited spatially concordant alternans, in which action potential duration alternates on a beat-to-beat basis. This alternation occurs concurrently with a beat-to-beat alternation in CV, such that the CV vs. BCL bifurcates at this alternans onset cycle length. We find that alternans onset is consistently at a shorter BCL for the Mix/Homogeneous-Polarized distribution, i.e., faster pacing is required to induce alternans. In Supporting Figure S5, we plot CV against the preceding diastolic interval (DI) (i.e., the CV restitution curve) for these conditions. We find that the CV restitution curve extends to shorter DI values for the Mix/Homogeneous-Polarized distribution, and in most cases, the curve is flatter, i.e., CV is less sensitive to changes in DI.

**Figure 12.**
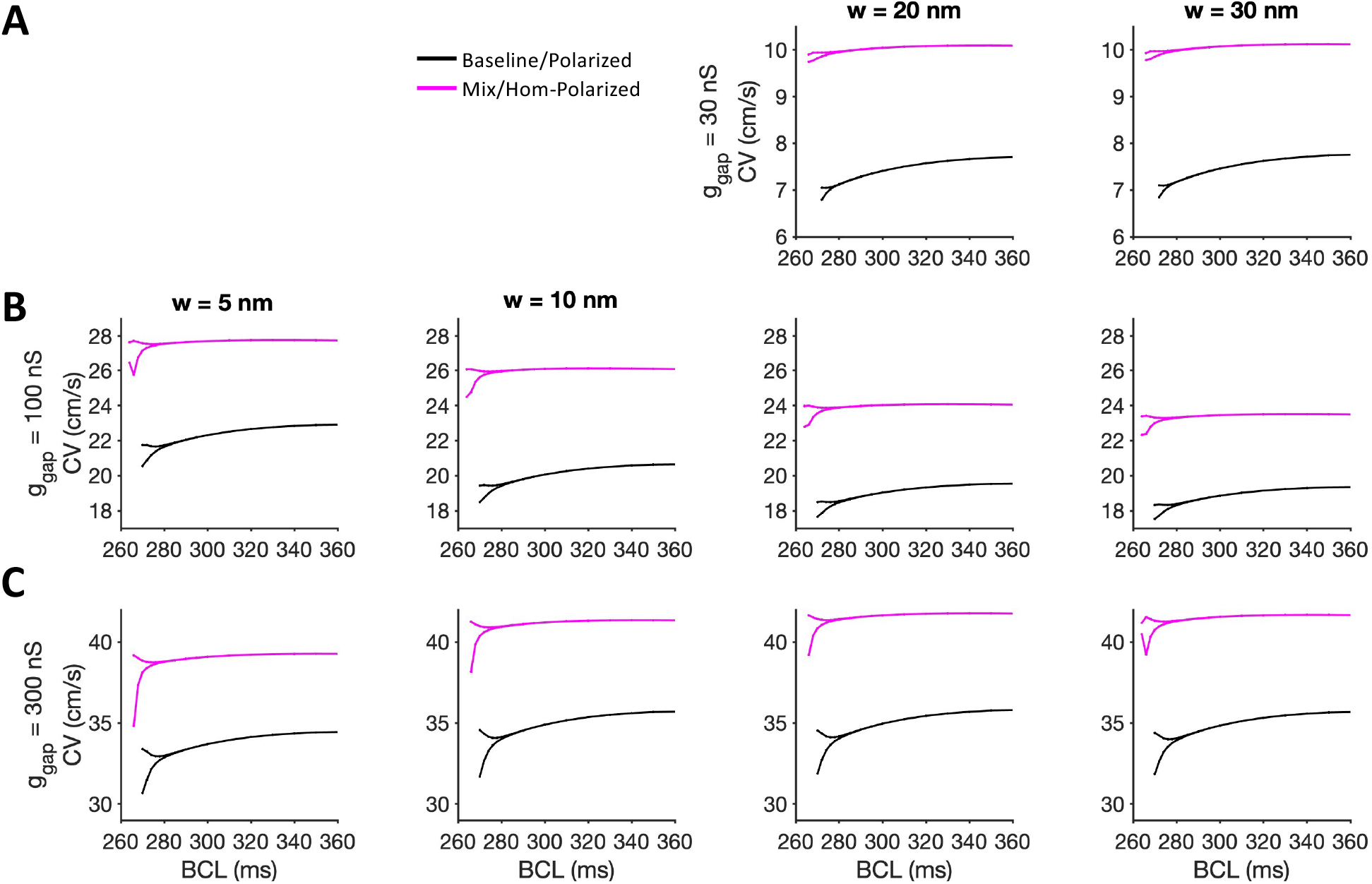
The Mix/Homogeneous-Polarized distribution exhibits faster conduction, compared with the Baseline/Polarized distribution, for all basic cycle lengths (BCLs). Conduction velocity (CV) is shown as a function of BCL for different values of cleft widths (w) and gap junction conductance g_gap_. Note, for the cases in the upper left panels (low g_gap_, narrow cleft width), conduction failed to capture at shorter BCL values.

Finally, we consider to what extent conduction differences due to the Mixed distribution depend on the shift in either the SSA or SSI curves. Thus, we consider cases in which the shifted I_Na_ subpopulation have only either a shift in the SSA or SSI relationships. In Figure 13A, we plot CV as function of g_gap_ for these two cases, with the Baseline/Polarized and Mix/Homogeneous-Polarized (with both SSA- and SSI-shifts) distributions shown for comparison. CV increases for increasing g_gap_, as in previous results. Importantly, for all g_gap_ and cleft widths, we find that the Mix/Homogeneous-Polarized distribution (magenta), with both SSA and SSI shifts, exhibits the largest CV, while the Baseline/Polarized distribution (black) exhibits the slowest CV. The Mix/Homogeneous-Polarized distribution, with only the SSA shift, has the next fastest CV, while the case with only the SSI shift is third.

**Figure 13.**
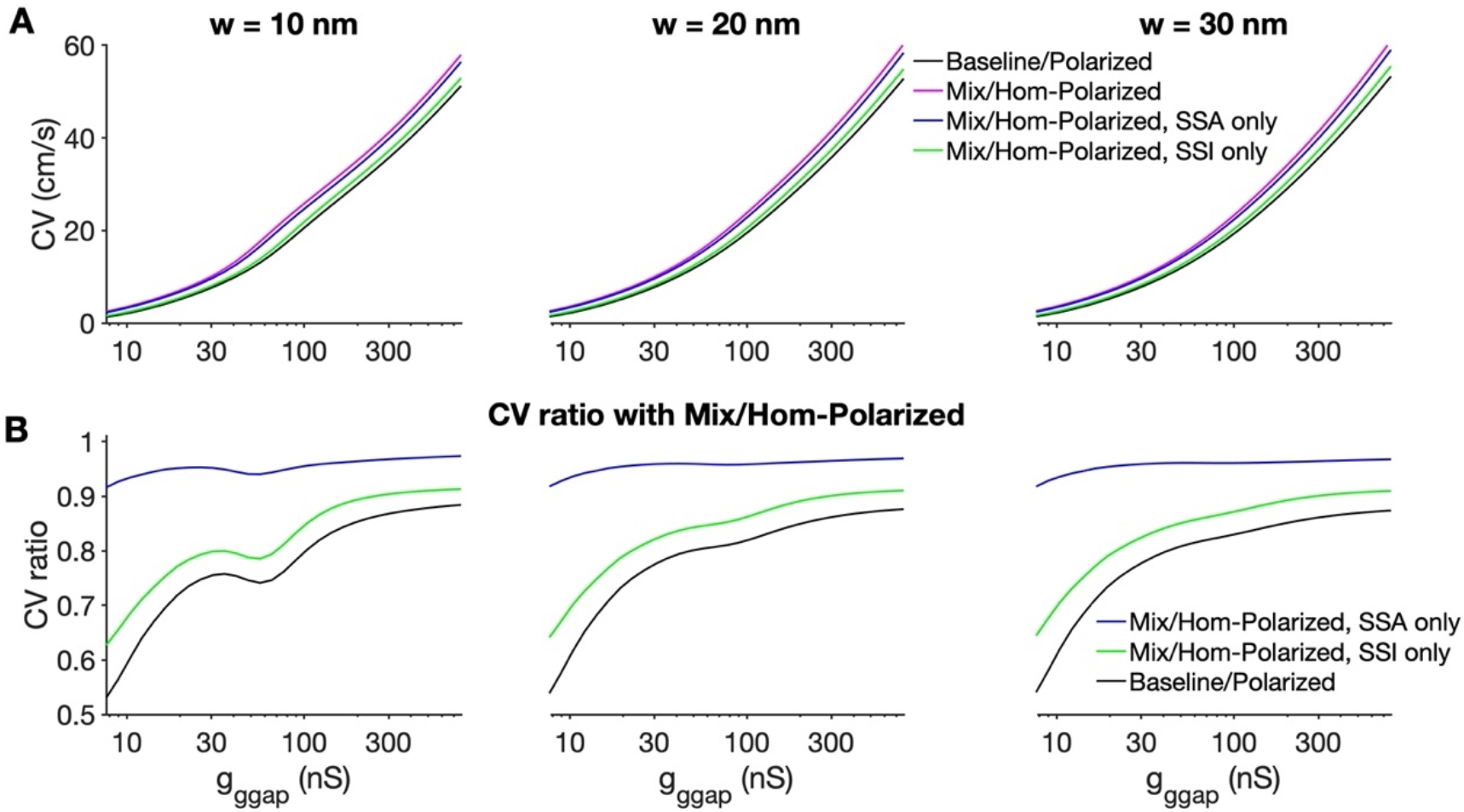
The shift in steady-state activation (SSA) predominantly reproduces the conduction changes in the shifted I_Na_ subpopulation. (A) Conduction velocity (CV) is shown as a function of gap junction conductance g_gap_ and cleft widths of 10 (right), 20 (middle), and 30 (right) nm, for the Baseline/Polarized (black) and Mix/Homogeneous-Polarized (both shifts, magenta; only SSA, blue; only SSI, green) distributions. (B) The ratio of CV values for different distributions, with the Mix/Homogeneous-Polarized distributions, are shown.

Relative to the Mix/Homogeneous-Polarized distribution, the CV ratio for tissue with only SSA-shifted I_Na_ is above 0.9, nearly independent of g_gap_ or cleft width (Figure 13B). In contrast, the CV ratio for the tissue with only SSI-shifted I_Na_ is consistently lower, varying between 0.6 and 0.9, generally decreasing as g_gap_ decreases. The CV ratio for the Baseline/Polarized distribution is the smallest and always less than 1 (Note: this curve is the inverse of the dashed line in Figure 6C). Thus, we find that, while both SSA and SSI shifts result in enhancement in conduction (relative to the Baseline/Polarized case), the shift in the SSA curve is primarily responsible for the increase in conduction observed in the Mix/Homogeneous-Polarized distribution.

## DISCUSSION

### Summary of main findings

In this study, we demonstrate that a subpopulation with distinct biophysical properties, specifically shifted SSI and SSA voltage-dependence, can modulate cardiac conduction. In a single cell, this subpopulation promotes an early AP upstroke, in a manner such that the shifted Na^+^ channels provide a proportionally greater contribution to the total Na^+^ charge during the upstroke. In a one-dimensional tissue model that accounts for the subcellular spatial localization of the different subpopulations, the shifted Na^+^ channel promote faster conduction. Further, the shifted Na^+^ channels result in greater CV sensitivity to changes in the intercellular cleft width, and this occurs across the entire range of physiological values for the gap junction conductance. The ID-localized shifted Na^+^ channels also provide a proportionally greater contribution to the total Na^+^ charge, in particular the post-junctional Na^+^ channels, while lateral membrane-localized Na^+^ channels contribute proportionally less, such that loss of the ID-localized Na^+^ channels result in greater conduction slowing. Additionally, we find that the ID-localized shifted Na^+^ channels enable conduction for faster pacing rates, in a manner that results in a flatter CV restitution curve. Finally, we demonstrate that the shift in the SSA curve is primarily responsible for the enhancement in conduction.

### Physiological sources and consequences of two Na^+^ channel subpopulations

We highlight that our computational study is agnostic as to the specific “source” for the different biophysical properties between the Na^+^ channel subpopulations, which could arise due to differences in interacting and scaffolding proteins at the distinct subcellular locations^1,2,8,56^. We speculate that association with different β subunits, which have been shown to alter Nav1.5 biophysical properties^5,6^, in different subcellular locations could also contribute to distinct Na^+^ subpopulations. Finally, there is increasing evidence for the presence of different Nav1.x isoforms in cardiac myocytes,^57,58^ which could also result in subpopulations with distinct biophysical properties. Our computational study provides a framework for investigating the potential impact of different Na^+^ channel subpopulations. Here, we specifically considered two subpopulations, with “baseline” and “shifted” biophysical properties based on measures from Lin et al^11^; we highlight that the specific subpopulation biophysical properties were based on one specific set of conditions. However, these properties can and likely do vary for different conditions and disease settings.

As a final investigation illustrating how different biophysical properties impact conduction, we measure CV for different values of 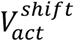 and 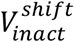 for the ID-localized Na^+^ channels in the Mix/Homogeneous-Polarized distribution (Figure 14). Across all cleft widths and gap junction conductances, a negative (left) shift in SSA and positive (right) shift in SSI both promote faster conduction, with shifts in SSA modulating CV with greater sensitivity. Thus, the directional shifts in both SSA and SSI curves in the physiological Na^+^ channel distribution both contribute to faster conduction. Further, we note that the large CV ranges observed due to shifts in the SSA and SSI curves of on the order of 10 mV. Thus, our study illustrates that biophysical properties differences can have important functional consequences for cardiac conduction and the contributions of the different subpopulation. Further, our approach could be naturally extended to consider multiple subpopulations with both distinct and mixed spatial localization, with biophysical properties based on interacting proteins, β subunits, and Nav isoforms. Future work will consider such complexities, in collaboration with experimental groups performing microscopy, patch clamp, and biophysical measurements.

**Figure 14.**
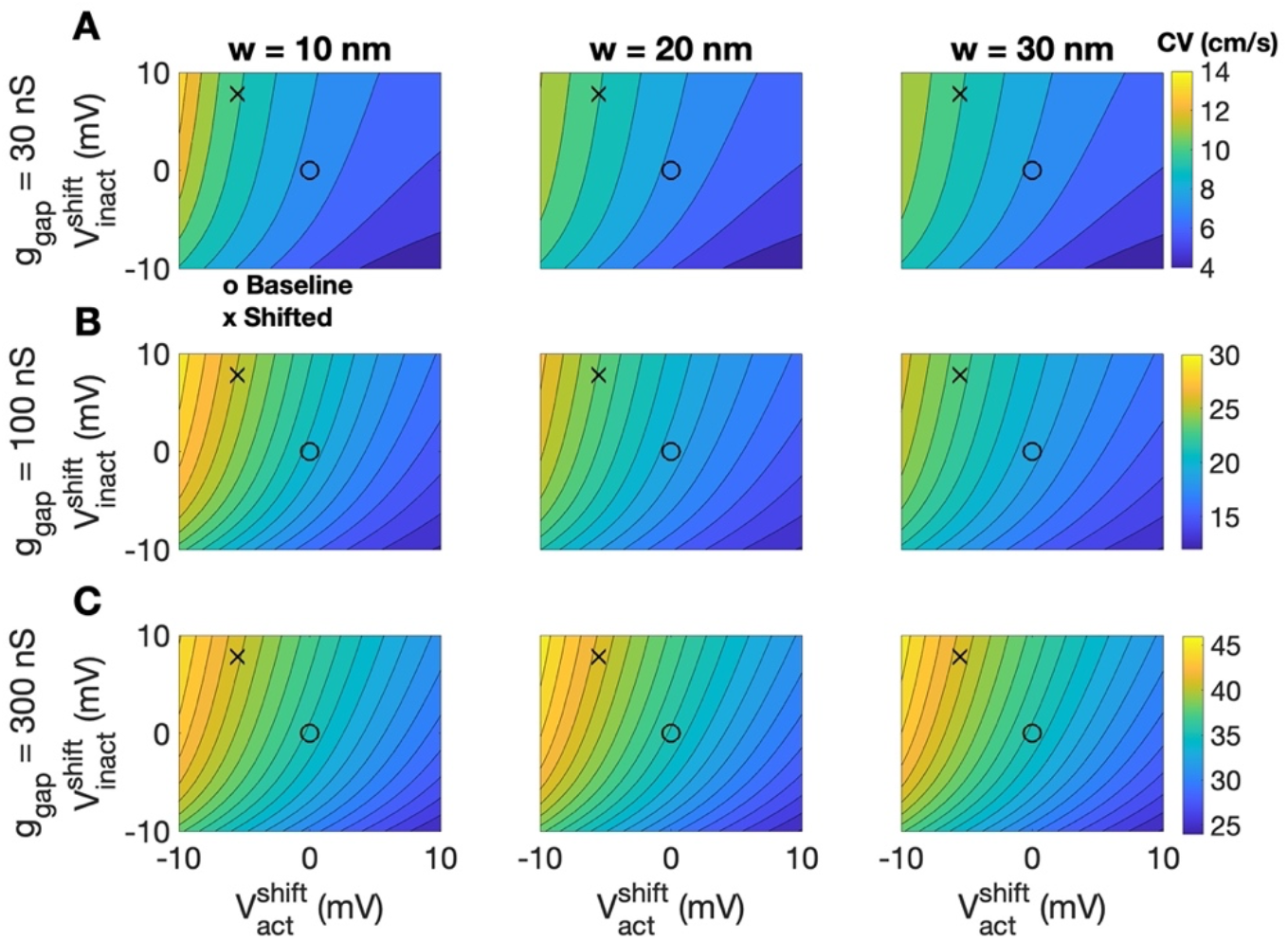
Negative shifts in steady-state activation and positive shifts in steady-state inactivation enhance conduction. Contour maps of the conduction velocity (CV) are shown as functions of 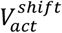 and 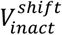, for different values of cleft widths (w) and gap junction conductance g_gap_, with 1-cm/s spacing. Parameters: Mix/Homogeneous-Polarized distribution, f_Na_ = 0.7.

We note that our computational study tests several predictions posed in the study from Lin and colleagues^11^. In their Discussion, the authors posit that Na^+^ channels in the middle of the cell “are mostly inactivated at a normal resting potential, leaving most of the burden of excitation to …[Na^+^] channels in the ID region.” Consideration of the Mix/Homogeneous-Polarized physiological Na^+^ distribution, our results mostly agree with this prediction (Figures 8-9). In particular, our results do predict that ID-localized Na^+^ channels provide the majority of the burden of excitation (quantified by the fraction of total Na^+^ charge), in particular the prejunctional ID Na^+^ channels. However, we predict that the lateral membrane-localized Na^+^ channels also contribute, albeit less than their respective proportion of the total Na^+^ conductance. Further, Lin and colleagues state that “[Middle Na^+^] channels have a minimum contribution to I_Na_ under control conditions but represent a functional reserve that can be upregulated by exogenous factors.” Our simulations also generally agree with these predictions. Interestingly, the lateral membrane-localized Na^+^ channels contribute the least to excitation as gap junctional coupling is reduced, while pre-junctional Na+ channels contribute the most for moderate values of gap junctional conductance (Figure 10). While our study does not specifically test the hypothesis that lateral membrane Na^+^ channels are upregulated by exogeneous factors under some conditions, our study does provide support for the notion of these channels serving as a functional reserve. That is, as f_Na_ decreases, the channels will provide a greater contribution to the burden of excitation. Additionally, while our study predicts that the loss of junctional Na^+^ channels results in greater conduction slowing, the loss of the lateral membrane-localized Na^+^ channels also slows conduction (Figures 11, S5), consistent with prior experimental studies^59^.

One interesting observation from the simulations across all 7 considered Na^+^ channel distributions is that the physiological distribution (i.e., the Mix/Homogeneous-Polarized), in general, did not support the fastest conduction (Figure 4); rather, the Shifted/Polarized distribution generally had the largest CV across different cleft widths and gap junction conductances. That is, the fastest conduction occurred for tissues with *only* the shifted Na^+^ channels, which naturally suggests the question of why the physiological distribution may be beneficial, compared with this Shifted/Polarized distribution of only shifted Na^+^ channels. First, our data demonstrates that the Mix/Homogeneous-Polarized distribution exhibits the greatest sensitivity in response to changes in cleft width (Figure 7). Thus, if we analogize conduction as being regulated by a series of “dials,” then gap junctional conductance is the “strongest” dial, highlighted by the wide range of CV values over the physiological range for g_gap_. While in this study, overall Na^+^ channel conductance is fixed, prior work has also shown strong sensitivity to this value,^25^ suggesting overall Na^+^ channel conductance is additional key “dial” modulating conduction. Our current study suggests that additional “dials” include the cleft width and Na^+^ channel biophysical properties and subcellular localization.

Additionally, we speculate that restricting Na^+^ channels with shifted biophysical properties to the ID and not the lateral membrane or t-tubules could potentially play an anti-arrhythmogenic protective role via several mechanisms. In the setting of dysfunctional calcium (Ca^2+^) handling, prior work has shown that spontaneous calcium release events can promote an influx of depolarizing current via the Na^+^/Ca^2+^ exchanger (NCX), which in turn can promote pro-arrhythmic delayed afterdepolarizations (DADs)^60–62^. Colocalization of shifted Na^+^ channels (specifically with left-shifted SSA) with the Ca^2+^ handling proteins in t-tubules would be more likely to activate due to the NCX current and thus drive DADs, such that restricting these shifted Na^+^ channels to the ID would be anti-arrhythmogenic. In contrast, due to inter-cellular cleft hyperpolarization described above, the subpopulation of shifted Na^+^ channels at the ID will, by design, activated earlier (Figures 8, S3) and is thus ideal for supporting robust conduction in concert with ephaptic mechanisms. Additionally, at faster pacing rates (and thus short DIs), prior studies have shown that steeper CV restitution curve promotes spatially discordant alternans, which can lead to conduction block and wavebreaks to initiate spiral waves and arrhythmias^63–65^, suggesting that the flatter CV restitution curve of the physiological distribution is also anti-arrhythmic (Figures 12, S6).

### Limitations, Future Directions, and Conclusions

We note potential limitations of our study. Our tissue simulations represent a one-dimensional “fiber” or chain of coupled cells that does not account for the complex and heterogeneous threedimensional geometry of the heart. This limits our ability to make predictions on the potential impact of Na^+^ channel subpopulations on transverse conduction or inherently higher dimensional electrical dynamics, such as spiral waves and arrhythmia initiation. Additionally, while we vary the distribution of Na^+^ channels and their respective biophysical properties, we do not consider the role of other ionic currents. While most other ionic currents will have minimal impact on conduction, the relative expression of the inward rectifier I_K1_ current has been shown to impact excitability via regulation of the resting membrane potential.^66^

Further, we assume that the biophysical properties and localization of the subpopulations are static; however, we speculate that these characteristics may be dynamic. Indeed, our study demonstrates that variability in these properties provide additional mechanisms for regulating conduction. Thus, our study can be considered a “snapshot” of conduction for a given Na^+^ channel distribution. We also do not consider additional degrees of heterogeneity, both within the tissue and within the ID. While to our knowledge, there are no measures of the spatial heterogeneity in Na^+^ channel distributions, it highly plausible that such heterogeneity exists within the ventricles, and we hypothesize that conduction is robust to some degree of spatial heterogeneity in these properties. This question is a focus of future work. Additionally, there is significant spatial heterogeneity within the ID, specifically within plicate and interplicate regions, in membrane structure and intermembrane separation. We have recently developed a finite-element framework to model ID structure and integrate the resulting heterogeneity in ID and cleft properties into a tissue-scale model.^26^ The broad parameter investigation performed here would have been computationally prohibitive with this more detailed model; however, we are also investigating the role of both ion channel organization and biophysical properties on conduction in future work.

In conclusion, we investigate the role of Na^+^ channel subpopulations with distinct biophysical properties and spatial localization on cardiac conduction. We find that ID-localized Na^+^ channels with shifted SSA and SSI biophysical properties support the majority of burden of excitation, and contribute to faster and more robust conduction, specifically sensitivity to tissue structural changes (i.e., cleft width), gap junctional coupling, and fast pacing rates. Our work supports the hypothesis that Na^+^ channel redistribution and biophysical properties regulation are critical mechanisms by which cardiac cells rapidly respond to perturbations to support electrical conduction.

## Acknowledgements.

This work was supported by the National Institutes of Health (NIH) R01HL138003.

## Data Availability

The simulation data is available upon reasonable request from the author.

## Notes

### Competing Interest Statement

The authors have declared no competing interest.

## References

1. Rivaud MR, Delmar M, Remme CA. Heritable arrhythmia syndromes associated with abnormal cardiac sodium channel function: ionic and non-ionic mechanisms. Cardiovasc Res. 2020;116(9):1557–1570. doi: 10.1093/cvr/cvaa082

2. Abriel H. Cardiac sodium channel Nav1.5 and interacting proteins: Physiology and pathophysiology. J Mol Cell Cardiol. 2010;48(1):2–11. doi: 10.1016/j.yjmcc.2009.08.025

3. Marchal GA, Remme CA. Subcellular diversity of Nav1.5 in cardiomyocytes: distinct functions, mechanisms and targets. J Physiol. Published online December 19, 2022:JP283086. doi: 10.1113/JP283086

4. Veeraraghavan R, Hoeker GS, Alvarez-Laviada A, et al. The adhesion function of the sodium channel beta subunit (β1) contributes to cardiac action potential propagation. eLife. 2018;7:e37610. doi: 10.7554/eLife.37610

5. Angsutararux P, Zhu W, Voelker TL, Silva JR. Molecular Pathology of Sodium Channel Beta-Subunit Variants. Front Pharmacol. 2021;12:761275. doi: 10.3389/fphar.2021.761275

6. Zhu W, Voelker TL, Varga Z, Schubert AR, Nerbonne JM, Silva JR. Mechanisms of noncovalent β subunit regulation of NaV channel gating. J Gen Physiol. 2017;149(8):813–831. doi: 10.1085/jgp.201711802

7. Watanabe H, Darbar D, Kaiser DW, et al. Mutations in Sodium Channel β1- and β2-Subunits Associated With Atrial Fibrillation. Circ Arrhythm Electrophysiol. 2009;2(3):268–275. doi: 10.1161/CIRCEP.108.779181

8. Shy D, Gillet L, Abriel H. Cardiac sodium channel NaV1.5 distribution in myocytes via interacting proteins: The multiple pool model. Biochim Biophys Acta BBA - Mol Cell Res. 2013;1833(4):886–894. doi: 10.1016/j.bbamcr.2012.10.026

9. Petitprez S, Zmoos AF, Ogrodnik J, et al. SAP97 and Dystrophin Macromolecular Complexes Determine Two Pools of Cardiac Sodium Channels Na v 1.5 in Cardiomyocytes. Circ Res. 2011;108(3):294–304. doi: 10.1161/CIRCRESAHA.110.228312

10. Balycheva M, Faggian G, Glukhov AV, Gorelik J. Microdomain–specific localization of functional ion channels in cardiomyocytes: an emerging concept of local regulation and remodelling. Biophys Rev. 2015;7(1):43–62. doi: 10.1007/s12551-014-0159-x

11. Lin X, Liu N, Lu J, et al. Subcellular heterogeneity of sodium current properties in adult cardiac ventricular myocytes. Heart Rhythm. 2011;8(12):1923–1930. doi: 10.1016/j.hrthm.2011.07.016

12. Veeraraghavan R, Lin J, Hoeker GS, Keener JP, Gourdie RG, Poelzing S. Sodium channels in the Cx43 gap junction perinexus may constitute a cardiac ephapse: an experimental and modeling study. Pflüg Arch - Eur J Physiol. 2015;467(10):2093–2105. doi: 10.1007/s00424-014-1675-z

13. Gillet L, Shy D, Abriel H. Elucidating sodium channel NaV1.5 clustering in cardiac myocytes using super-resolution techniques. Cardiovasc Res. 2014;104(2):231–233. doi: 10.1093/cvr/cvu221

14. Agullo-Pascual E, Lin X, Leo-Macias A, et al. Super-resolution imaging reveals that loss of the C-terminus of connexin43 limits microtubule plus-end capture and NaV1.5 localization at the intercalated disc. Cardiovasc Res. 2014;104(2):371–381. doi: 10.1093/cvr/cvu195

15. Marchal GA, Jouni M, Chiang DY, et al. Targeting the Microtubule EB1-CLASP2 Complex Modulates Na v 1.5 at Intercalated Discs. Circ Res. 2021;129(3):349–365. doi: 10.1161/CIRCRESAHA.120.318643

16. Kucera JP, Rohr S, Rudy Y. Localization of Sodium Channels in Intercalated Disks Modulates Cardiac Conduction. Circ Res. 2002;91(12):1176–1182. doi: 10.1161/01.RES.0000046237.54156.0A

17. Mori Y, Fishman GI, Peskin CS. Ephaptic conduction in a cardiac strand model with 3D electrodiffusion. Proc Natl Acad Sci U S A. 2008;105(17):6463–6468. doi: 10.1073/pnas.0801089105

18. Lin J, Keener JP. Modeling electrical activity of myocardial cells incorporating the effects of ephaptic coupling. Proc Natl Acad Sci U S A. 2010;107(49):20935–20940. doi: 10.1073/pnas.1010154107

19. Rhett JM, Veeraraghavan R, Poelzing S, Gourdie RG. The perinexus: Sign-post on the path to a new model of cardiac conduction? Trends Cardiovasc Med. 2013;23(6):222–228. doi: 10.1016/j.tcm.2012.12.005

20. Anastassiou CA, Perin R, Markram H, Koch C. Ephaptic coupling of cortical neurons. Nat Neurosci. 2011;14(2):217–223. doi: 10.1038/nn.2727

21. Sperelakis N. The possibility of propagation between myocardial cells not connected by low-resistance pathways. Adv Exp Med Biol. 1983;161:1–23. doi: 10.1007/978-1-4684-4472-8_1

22. Sperelakis N, Mann JE. Evaluation of electric field changes in the cleft between excitable cells. J Theor Biol. 1977;64(1):71–96. doi: 10.1016/0022-5193(77)90114-x

23. Sperelakis N, McConnell K. Electric field interactions between closely abutting excitable cells. IEEE Eng Med Biol Mag Q Mag Eng Med Biol Soc. 2002;21(1):77–89. doi: 10.1109/51.993199

24. Pertsov AM, Medvinskiĭ AB. Electric coupling in cells without highly permeable cell contacts. Biofizika. 1976;21(4):698–670.

25. Nowak MB, Veeraraghavan R, Poelzing S, Weinberg SH. Cellular Size, Gap Junctions, and Sodium Channel Properties Govern Developmental Changes in Cardiac Conduction. Front Physiol. 2021;12:731025. doi: 10.3389/fphys.2021.731025

26. Moise N, Struckman HL, Dagher C, Veeraraghavan R, Weinberg SH. Intercalated disk nanoscale structure regulates cardiac conduction. J Gen Physiol. 2021;153(8):e202112897. doi: 10.1085/jgp.202112897

27. Weinberg SH. Ephaptic coupling rescues conduction failure in weakly coupled cardiac tissue with voltage-gated gap junctions. Chaos Interdiscip J Nonlinear Sci. 2017;27(9):093908. doi: 10.1063/1.4999602

28. Nowak MB, Poelzing S, Weinberg SH. Mechanisms underlying age-associated manifestation of cardiac sodium channel gain-of-function. J Mol Cell Cardiol. 2021;153:60–71. doi: 10.1016/j.yjmcc.2020.12.008

29. Nowak MB, Greer-Short A, Wan X, et al. Intercellular Sodium Regulates Repolarization in Cardiac Tissue with Sodium Channel Gain of Function. Biophys J. 2020;118(11):2829–2843. doi: 10.1016/j.bpj.2020.04.014

30. Greer-Short A, George SA, Poelzing S, Weinberg SH. Revealing the Concealed Nature of Long-QT Type 3 Syndrome. Circ Arrhythm Electrophysiol. 2017;10(2). doi: 10.1161/CIRCEP.116.004400

31. Yu JK, Liang JA, Weinberg SH, Trayanova NA. Computational modeling of aberrant electrical activity following remuscularization with intramyocardially injected pluripotent stem cell-derived cardiomyocytes. J Mol Cell Cardiol. 2022;162:97–109. doi: 10.1016/j.yjmcc.2021.08.011

32. Veeraraghavan R, Lin J, Keener JP, Gourdie R, Poelzing S. Potassium channels in the Cx43 gap junction perinexus modulate ephaptic coupling: an experimental and modeling study. Pflugers Arch. 2016;468(10):1651–1661. doi: 10.1007/s00424-016-1861-2

33. Lin J, Keener JP. Ephaptic coupling in cardiac myocytes. IEEE Trans Biomed Eng. 2013;60(2):576–582. doi: 10.1109/TBME.2012.2226720

34. Lin J, Keener JP. Microdomain effects on transverse cardiac propagation. Biophys J. 2014;106(4):925–931. doi: 10.1016/j.bpj.2013.11.1117

35. Ly C, Weinberg SH. Automaticity in ventricular myocyte cell pairs with ephaptic and gap junction coupling. Chaos Interdiscip J Nonlinear Sci. 2022;32(3):033123. doi: 10.1063/5.0085291

36. Poelzing S, Weinberg SH, Keener JP. Initiation and entrainment of multicellular automaticity via diffusion limited extracellular domains. Biophys J. 2021;120(23):5279–5294. doi: 10.1016/j.bpj.2021.10.034

37. Hand PE, Peskin CS. Homogenization of an electrophysiological model for a strand of cardiac myocytes with gap-junctional and electric-field coupling. Bull Math Biol. 2010;72(6):1408–1424. doi: 10.1007/s11538-009-9499-2

38. Wei N, Mori Y, Tolkacheva EG. The dual effect of ephaptic coupling on cardiac conduction with heterogeneous expression of connexin 43. J Theor Biol. 2016;397:103–114. doi: 10.1016/j.jtbi.2016.02.029

39. Wei N, Tolkacheva EG. Mechanisms of arrhythmia termination during acute myocardial ischemia: Role of ephaptic coupling and complex geometry of border zone. PloS One. 2022;17(3):e0264570. doi: 10.1371/journal.pone.0264570

40. Hichri E, Abriel H, Kucera JP. Distribution of cardiac sodium channels in clusters potentiates ephaptic interactions in the intercalated disc: Ephaptic effects in intercalated discs. J Physiol. 2018;596(4):563–589. doi: 10.1113/JP275351

41. Ivanovic E, Kucera JP. Tortuous Cardiac Intercalated Discs Modulate Ephaptic Coupling. Cells. 2022;11(21):3477. doi: 10.3390/cells11213477

42. Ivanovic E, Kucera JP. Localization of Na ^+^ channel clusters in narrowed perinexi of gap junctions enhances cardiac impulse transmission via ephaptic coupling: a model study. J Physiol. 2021;599(21):4779–4811. doi: 10.1113/JP282105

43. Luo CH, Rudy Y. A model of the ventricular cardiac action potential. Depolarization, repolarization, and their interaction. Circ Res. 1991;68(6):1501–1526. doi: 10.1161/01.res.68.6.1501

44. Wu X, Hoeker GS, Blair GA, et al. Hypernatremia and intercalated disc edema synergistically exacerbate long-QT syndrome type 3 phenotype. Am J Physiol-Heart Circ Physiol. 2021;321(6):H1042–H1055. doi: 10.1152/ajpheart.00366.2021

45. Desplantez T, Dupont E, Severs NJ, Weingart R. Gap Junction Channels and Cardiac Impulse Propagation. J Membr Biol. 2007;218(1-3):13–28. doi: 10.1007/s00232-007-9046-8

46. Kwak BrendaR, Jongsma HaboJ. Regulation of cardiac gap junction channel permeability and conductance by several phosphorylating conditions. Mol Cell Biochem. 1996;157(1-2). doi: 10.1007/BF00227885

47. McCain ML, Desplantez T, Geisse NA, et al. Cell-to-cell coupling in engineered pairs of rat ventricular cardiomyocytes: relation between Cx43 immunofluorescence and intercellular electrical conductance. Am J Physiol-Heart Circ Physiol. 2012;302(2):H443–H450. doi: 10.1152/ajpheart.01218.2010

48. Moreno AP, Rook MB, Fishman GI, Spray DC. Gap junction channels: distinct voltage-sensitive and -insensitive conductance states. Biophys J. 1994;67(1):113–119. doi: 10.1016/S0006-3495(94)80460-6

49. Valiunas V, Beyer EC, Brink PR. Cardiac Gap Junction Channels Show Quantitative Differences in Selectivity. Circ Res. 2002;91(2):104–111. doi: 10.1161/01.RES.0000025638.24255.AA

50. Verheule S, van Kempen MJA, Welscher PHJA te, Kwak BR, Jongsma HJ. Characterization of Gap Junction Channels in Adult Rabbit Atrial and Ventricular Myocardium. Circ Res. 1997;80(5):673–681. doi: 10.1161/01.RES.80.5.673

51. White RL, Doeller JE, Verselis VK, Wittenberg BA. Gap junctional conductance between pairs of ventricular myocytes is modulated synergistically by H+ and Ca++. J Gen Physiol. 1990;95(6):1061–1075. doi: 10.1085/jgp.95.6.1061

52. Nielsen MS, Nygaard Axelsen L, Sorgen PL, Verma V, Delmar M, Holstein-Rathlou N. Gap Junctions. In: Terjung R, ed. Comprehensive Physiology. 1st ed. Wiley; 2012:1981–2035. doi: 10.1002/cphy.c110051

53. Rudisuli, A A, Weingart R. Electrical properties of gap junction channels in guinea-pig ventricular cell pairs revealed by exposure to heptanol. Pfl⍰gers Arch Eur J Physiol. 1989;415(1):12–21. doi: 10.1007/BF00373136

54. Weingart R. Electrical properties of the nexal membrane studied in rat ventricular cell pairs. J Physiol. 1986;370(1):267–284. doi: 10.1113/jphysiol.1986.sp015934

55. Wittenberg BA, White RL, Ginzberg RD, Spray DC. Effect of calcium on the dissociation of the mature rat heart into individual and paired myocytes: electrical properties of cell pairs. Circ Res. 1986;59(2):143–150. doi: 10.1161/01.RES.59.2.143

56. Rivaud MR, Agullo-Pascual E, Lin X, et al. Sodium Channel Remodeling in Subcellular Microdomains of Murine Failing Cardiomyocytes. J Am Heart Assoc. 2017;6(12):e007622. doi: 10.1161/JAHA.117.007622

57. Tarasov M, Struckman HL, Olgar Y, et al. NaV1.6 dysregulation within myocardial T-tubules by D96V calmodulin enhances proarrhythmic sodium and calcium mishandling. J Clin Invest. Published online February 23, 2023:e152071. doi: 10.1172/JCI152071

58. Westenbroek RE, Bischoff S, Fu Y, Maier SKG, Catterall WA, Scheuer T. Localization of sodium channel subtypes in mouse ventricular myocytes using quantitative immunocytochemistry. J Mol Cell Cardiol. 2013;64:69–78. doi: 10.1016/j.yjmcc.2013.08.004

59. Shy D, Gillet L, Ogrodnik J, et al. PDZ domain-binding motif regulates cardiomyocyte compartment-specific NaV1.5 channel expression and function. Circulation. 2014;130(2):147–160. doi: 10.1161/CIRCULATIONAHA.113.007852

60. Bers DM, Despa S, Bossuyt J. Regulation of Ca2+ and Na+ in normal and failing cardiac myocytes. Ann N Y Acad Sci. 2006;1080:165–177. doi: 10.1196/annals.1380.015

61. Bögeholz N, Pauls P, Bauer BK, et al. Suppression of Early and Late Afterdepolarizations by Heterozygous Knockout of the Na+/Ca2+ Exchanger in a Murine Model. Circ Arrhythm Electrophysiol. 2015;8(5):1210–1218. doi: 10.1161/CIRCEP.115.002927

62. Bögeholz N, Pauls P, Kaese S, et al. Triggered activity in atrial myocytes is influenced by Na+/Ca2+ exchanger activity in genetically altered mice. J Mol Cell Cardiol. 2016;101:106–115. doi: 10.1016/j.yjmcc.2016.11.004

63. Qu Z, Garfinkel A, Chen PS, Weiss JN. Mechanisms of discordant alternans and induction of reentry in simulated cardiac tissue. Circulation. 2000;102(14):1664–1670. doi: 10.1161/01.cir.102.14.1664

64. Weiss JN, Karma A, Shiferaw Y, Chen PS, Garfinkel A, Qu Z. From pulsus to pulseless: the saga of cardiac alternans. Circ Res. 2006;98(10):1244–1253. doi: 10.1161/01.RES.0000224540.97431.f0

65. Choi BR, Jang W, Salama G. Spatially discordant voltage alternans cause wavebreaks in ventricular fibrillation. Heart Rhythm. 2007;4(8):1057–1068. doi: 10.1016/j.hrthm.2007.03.037

66. Dhamoon AS, Jalife J. The inward rectifier current (IK1) controls cardiac excitability and is involved in arrhythmogenesis. Heart Rhythm. 2005;2(3):316–324. doi: 10.1016/j.hrthm.2004.11.012

