## Supplemental Figures for "Sodium channel subpopulations with distinct biophysical properties and subcellular localization enhance cardiac conduction"

### SUPPORTING MATERIAL

#### Supplemental Figures.

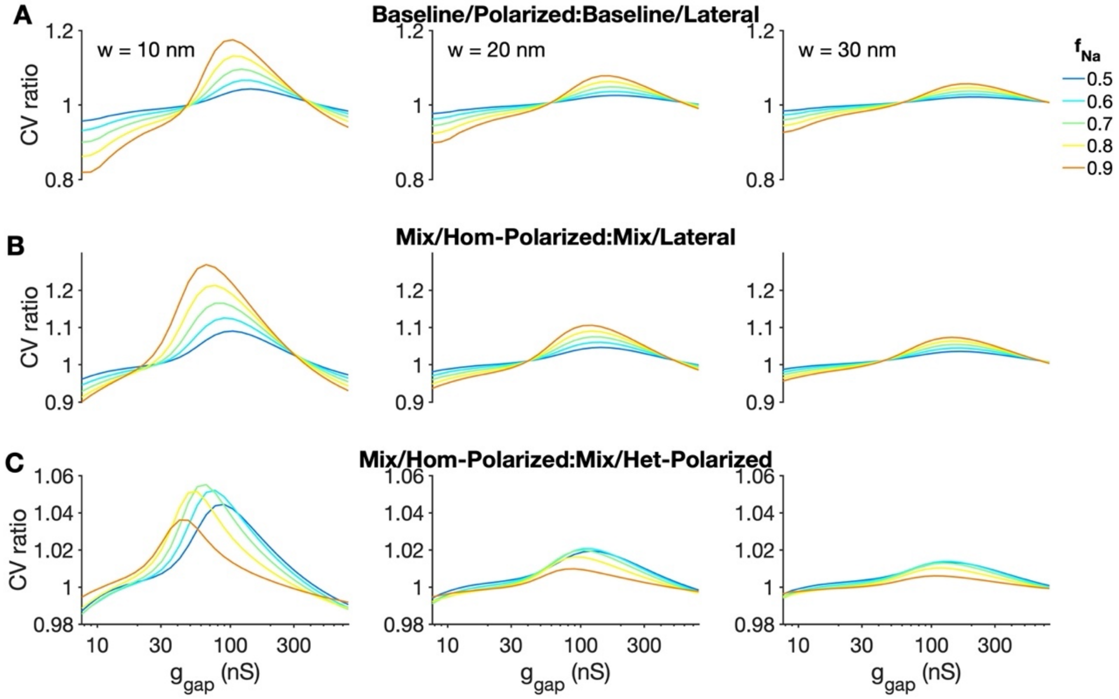


Figure S1. Larger fractions of shifted $I_{Na}$ and polarized $I_{Na}$ result greater dependence on $Na^+$ channel distribution. (A-C) The ratio of CV values for different $Na^+$ channel distribution combinations are shown as a function of gap junction conductance $g_{gap}$, for cleft widths of 10 (left), 20 (middle), and 30 (right) nm and different values of $f_{Na}$.

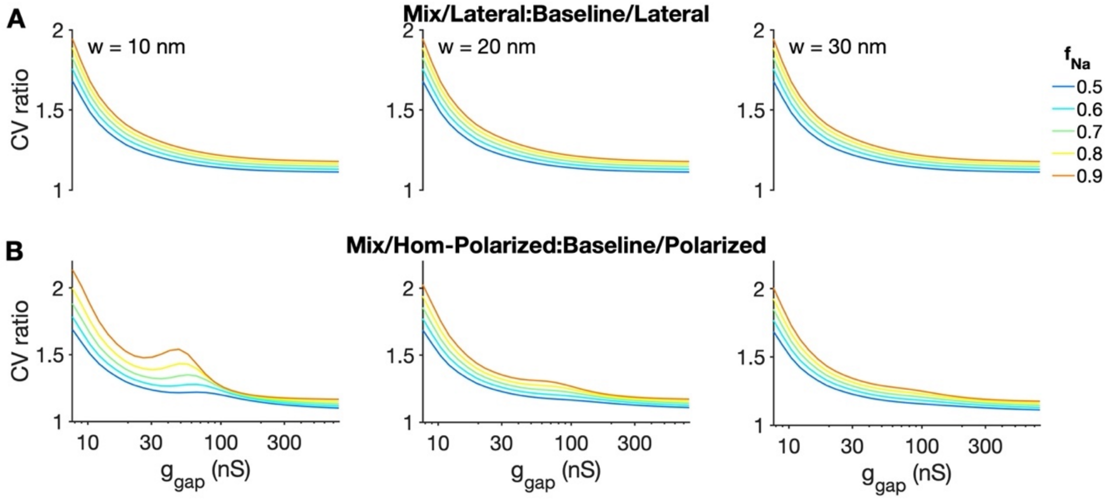


Figure S2. Larger fractions of shifted $I_{Na}$ and polarized $I_{Na}$ result greater dependence on $Na^+$ channel distribution. (A-C) The ratio of CV values for different $Na^+$ channel distribution combinations are shown as a function of gap junction conductance $g_{gap}$, for cleft widths of 10 (left), 20 (middle), and 30 (right) nm and different values of $f_{Na}$.

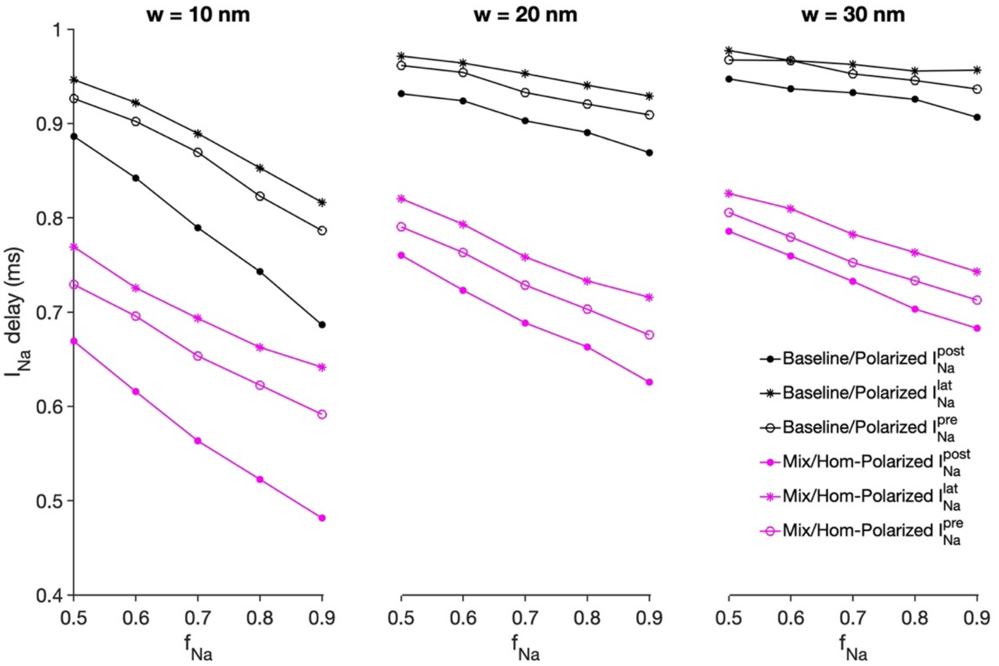


Figure S3. Post-junctional and lateral $I_{Na}$ activate earlier for narrow clefts and the Mix/Homogeneous-Polarized distribution. The delay in timing between the $I_{Na}$ peaks (on cell 25) and the cell 24 pre-junctional membrane is shown as a function of $f_{Na}$ for the Baseline/Polarized (black) and Mix/Homogeneous-Polarized (magenta) distributions. Parameters: $g_{gap}$ = 100 nS.

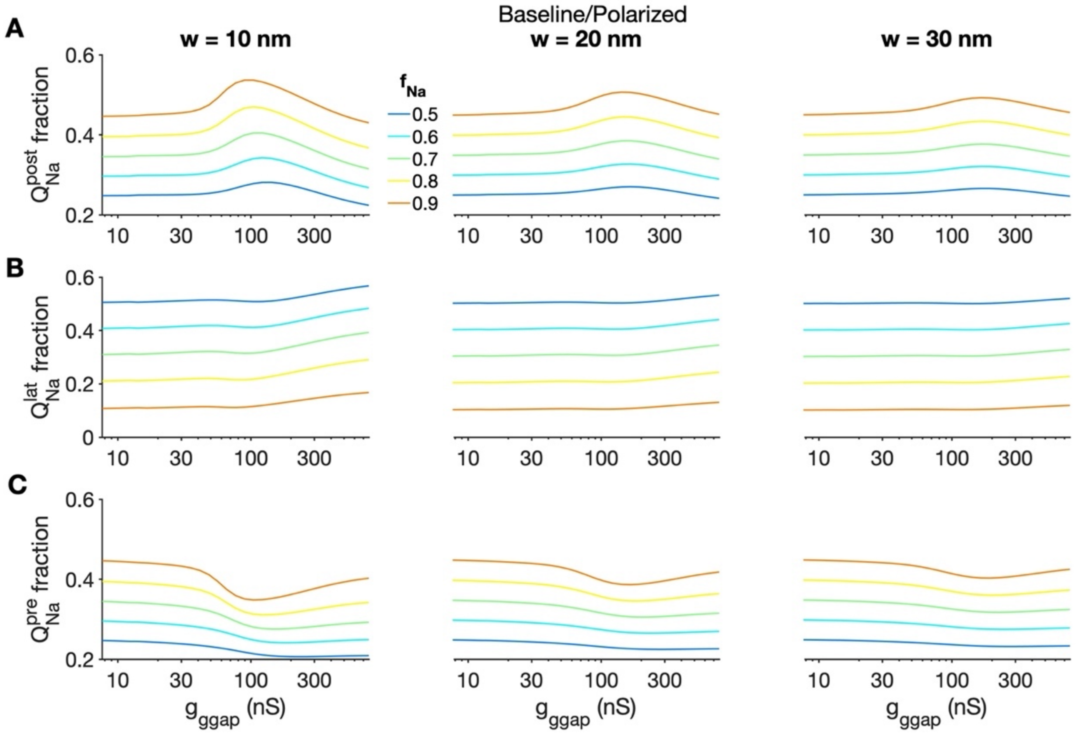


Figure S4. Post-junctional $I_{Na}$ contribution is enhanced for moderate gap junction coupling. The fraction of $Na^+$ charge $Q_{Na}$ carried by (A) post-junctional, (B) lateral, and (C) pre-junctional $I_{Na}$ is shown as a function of gap junction conductance $g_{gap}$ for cleft widths of 10 (left), 20 (middle), and 30 (right) nm and different values of $f_{Na}$ for the Baseline/Polarized distribution.

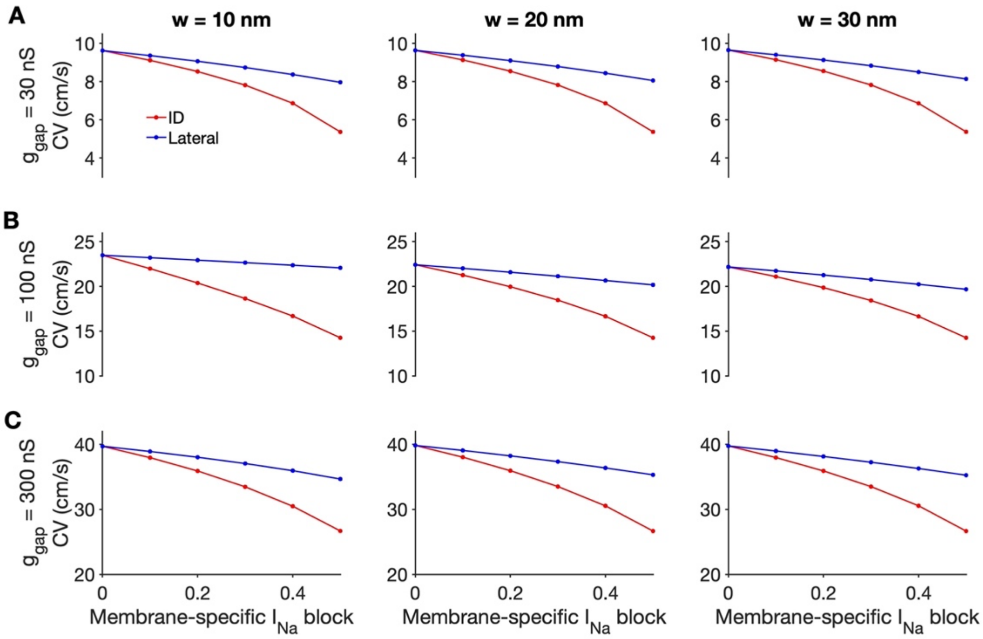


Figure S5. Membrane-specific $I_{Na}$ block in the Mix/Homogeneous-Polarized distribution. (A-C) Conduction velocity (CV) is shown as a function of the $I_{Na}$ conductance block of the ID (red) or lateral (blue) membrane, for different values of cleft width w and gap junction conductance $g_{gap}$. Parameters: $f_{Na}$ = 0.5.

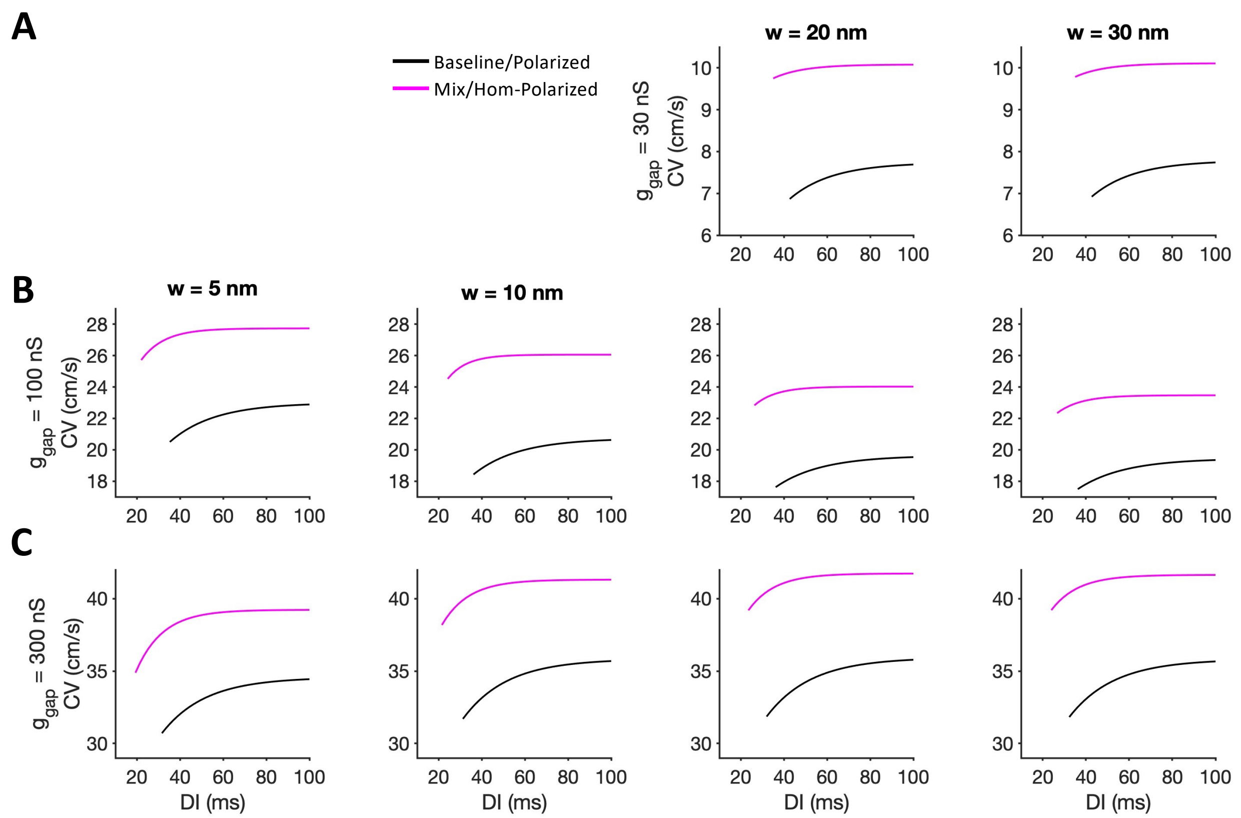


Figure S6. The Mix/Homogeneous-Polarized distribution exhibits faster conduction, compared with the Baseline/Polarized distribution, for all diastolic intervals (Dis). Conduction velocity (CV) is shown as a function of DI for different cleft widths w and gap junction conductance $g_{gap}$. For most conditions, the Mix/Homogeneous-Polarized distribution (magenta) CV restitution curve is flatter, compared with the Baseline/Polarized distribution (black). Note, for the cases in the upper left panels (low $g_{gap}$, narrow cleft width), conduction failed to capture at shorter BCL values.
